# Infant acquisition of Stickland-metabolizing bacteria prevents *Clostridium botulinum* infection

**DOI:** 10.64898/2026.06.23.734129

**Authors:** Nobuhide Kobayashi, Yotaro Kodaira, Jiayue Yang, Takuhiro Matsumura, Aki Yamaguchi, Yuta Arai, Daisuke Takahashi, Hiroki Toriumi, Seiga Komiyama, Kanon Iwata, Natsuki Haga, Yukina Nishida, Kazuki Saito, Daisuke Motooka, Yuki Matsumoto, Shota Nakamura, Taizo Wada, Shinji Fukuda, Koji Hase, Yukako Fujinaga

**Affiliations:** Department of Bacteriology, Graduate School of Medical Sciences, Kanazawa University, Kanazawa, Ishikawa 920-8640, Japan; Division of Biochemistry, Faculty of Pharmacy, Keio University, Tokyo 105-8512, Japan; Institute for Advanced Biosciences, Keio University, Tsuruoka, Yamagata 997-0052, Japan; Immunosurveillance Laboratory, The Francis Crick Institute, London, UK; Department of Infection Metagenomics, Bioinformatics Center, Research Institute for Microbial Diseases (RIMD), The University of Osaka, Suita, Osaka, 565-0871, Japan; Integrated Frontier Research for Medical Science Division, Institute for Open and Transdisciplinary Research Initiatives, The University of Osaka, Suita, Osaka, 565-0871, Japan; Center for Infectious Disease Education and Research, The University of Osaka, Suita, Osaka, 565-0871, Japan; Department of Pediatrics, Kanazawa University Hospital, Kanazawa, Ishikawa 920-8640, Japan; Innovative Microbiome Therapy Research Center, Juntendo University Graduate School of Medicine, Tokyo 113-8421, Japan; Gut Environmental Design Group, Kanagawa Institute of Industrial Science and Technology, Kawasaki, Kanagawa 210-0821, Japan; Transborder Medical Research Center, Institute of Medicine, University of Tsukuba, Tsukuba, Ibaraki 305-8575, Japan; Division of Commensal Microbiology, The Institute of Medical Science, The University of Tokyo (IMSUT), Tokyo 108-8639, Japan; International Vaccine Design Center, The Institute of Medical Science, The University of Tokyo (IMSUT), Tokyo 108-8639, Japan; Institute of Fermentation Sciences (IFeS), Faculty of Food and Agricultural Sciences, Fukushima University, Fukushima 960-1296, Japan

## Abstract

Infant botulism is caused by intestinal colonization with *Clostridium botulinum*, whereas healthy adults are resistant. Although the gut microbiota has long been implicated in protection against *C. botulinum*, its bacterial basis has remained unknown. Here, we show that specific amino acid-metabolizing bacteria confer resistance to *C. botulinum* colonization through nutrient competition. Longitudinally collected human infant microbiotas exhibited a transition from susceptibility to resistance after transplantation into germ-free mice. 5-Aminovalerate marked the resistant microbiota, implicating Stickland metabolism. Resistant microbiotas were enriched in Stickland-metabolizing Clostridia, including *Clostridioides difficile*. Metabolomics revealed overlapping amino acid utilization, and *C. difficile* suppressed *C. botulinum* expansion through amino acid competition. These findings demonstrate that acquisition of Stickland-metabolizing Clostridia during infancy prevents *C. botulinum* infection through competition for shared amino-acid-dependent nutritional niches.

## Main Text

Infant botulism is caused mainly by group I (proteolytic) *Clostridium botulinum*, and more rarely by botulinum neurotoxin (BoNT)-producing strains of *C. butyricum* and *C. baratii* (*1*). When infants under 1 year of age ingest spores of these bacteria, the spores germinate in the anaerobic environment of the lower intestine and subsequently produce BoNT, causing severe paralysis (*1*). Although human botulinum antitoxin is available for treatment (*2*), no strategy currently exists to prevent *C. botulinum* infection. Healthy older children and adults are generally resistant, whereas adults receiving high doses of antibiotics or having intestinal disorders occasionally become infected with *C. botulinum* and develop adult intestinal botulism (*3*). Early studies in the 1980s using rodent models revealed that conventional adult mice and rats were resistant to oral infection with *C. botulinum*, whereas germ-free (GF) or antibiotic-treated mice were susceptible (*4–7*). Furthermore, acquisition of an adult gut microbiota conferred resistance to *C. botulinum* infection in GF mice (*4*). A subsequent study demonstrated that mice harboring a defined intestinal microbiota exhibited resistance to *C. botulinum* colonization (*8*), suggesting that specific microbial components mediate protection, although the responsible bacteria remained unidentified. Together, these studies established the gut microbiota as a key determinant of resistance to *C. botulinum* infection. However, despite these pioneering observations, a longstanding question has remained unresolved since infant botulism was first medically recognized in 1976: which members of the gut microbiota confer resistance to *C. botulinum*, and by what mechanisms? Addressing this question may enable microbiota-based prevention of infant botulism.

Maturation of the infant gut microbiota is thought to underlie the acquisition of resistance to intestinal pathogens, yet the underlying mechanisms remain poorly understood (*9–11*). In mice, the neonatal gut microbiota is highly susceptible to infection by *Salmonella enterica* serovar Typhimurium and *Citrobacter rodentium*, whereas colonization by Clostridiales, a bacterial group abundant in adults, confers colonization resistance (CR) against these enteropathogenic bacteria through unknown mechanisms (*12*). However, how maturation of the infant gut microbiota contributes to CR against individual pathogens remains, particularly in humans, largely unclear.

## Results

### Commensal Clostridia consortia prevent *C. botulinum* infection

Microbiota-mediated CR against pathogenic bacteria is broadly classified into 2 categories: direct CR mediated through bacterial interactions, and indirect CR mediated by the host immune responses (*13*). We first investigated whether host immunity contributes to CR, because *C. botulinum* potentially induces inflammatory responses (*14*). The following experiments were performed using *C. botulinum* type A 62A, a group I (proteolytic) strain previously employed in infection models (*4*, *6–8*, *15*, *16*). Proteomic analyses demonstrated induction of both mucosal and systemic immune responses following infection (fig. S1A–C). To clarify the role of the host immune system, we compared susceptibility to *C. botulinum* infection in *Myd88*^−/−^*Trif*^−/−^ mice lacking Toll-like receptor signaling (*17*) and *Rag2*^−/−^*Jak3*^−/−^ mice lacking adaptive lymphocytes and natural killer (NK) cells (fig. S1D) (*18*). Both immunodeficient strains became susceptible following antibiotic treatment, whereas untreated mice remained resistant (fig. S1E–J). These results indicate that direct CR, rather than host immunity, is the primary protective mechanism against *C. botulinum* infection.

Previous epidemiological studies in human infants have suggested an association between microbiota composition and susceptibility to infant botulism(*19–21*); however, the bacterial taxa responsible for CR acquisition remain unknown. In experimental mouse models, conventional ICR mice are susceptible to *C. botulinum* infection at 7–13 days of age (*15*). To investigate the relationship between microbiota maturation and resistance acquisition while minimizing the confounding effects of host physical development, we transplanted cecal contents from infant specific pathogen-free (SPF) mice (4-, 10-, or 18-day-old) into age-matched adult GF mice and compared susceptibility to *C. botulinum* infection 1 week after transplantation (Fig. 1A). 16S ribosomal RNA (rRNA) gene amplicon sequencing revealed that *Lactobacillaceae* and *Enterococcaceae* were consistently abundant in mice transplanted with microbiota from 4- and 10-day-old donors, whereas the bacterial classes Clostridia and Bacteroidia predominated in mice transplanted with 18-day-old microbiota but were rarely detected in the other groups (Fig. 1B). Following infection, mice transplanted with microbiota from 4-or 10-day-old donors permitted robust colonization by *C. botulinum* and developed severe botulism symptoms, similar to GF mice (Fig. 1C–E). Notably, microbiota derived from 4-day-old mice failed to confer resistance despite being obtained from mice younger than the previously reported susceptible period (*15*), suggesting that resistance in neonatal mice may be mediated primarily by host factors rather than the gut microbiota. In contrast, mice transplanted with microbiota from 18-day-old donors exhibited resistance, with no detectable *C. botulinum* colonization or botulism symptoms throughout the observation period. These findings suggested that Clostridia and Bacteroidia are characteristic components of the resistant microbiota. To further identify bacterial groups associated with CR, we treated adult SPF mice with different antibiotics to selectively perturb distinct components of the gut microbiota (Fig. 1F, G). Mice treated with ampicillin, vancomycin, or metronidazole became susceptible to infection, exhibiting both intestinal colonization by *C. botulinum* and severe botulism symptoms. In contrast, mice treated with polymyxin B or erythromycin remained resistant, similar to vehicle-treated controls, with neither detectable colonization nor disease development (Fig. 1H–J). Indicator-species analysis comparing susceptible and resistant microbiota revealed that taxa associated with the resistant microbiota were predominantly members of the class Clostridia (Fig. 1K), which accounted for 38 of 43 resistant-associated indicator genera (table S1). These results identify Clostridia as a key bacterial group associated with CR against *C. botulinum*.

**Fig. 1.**
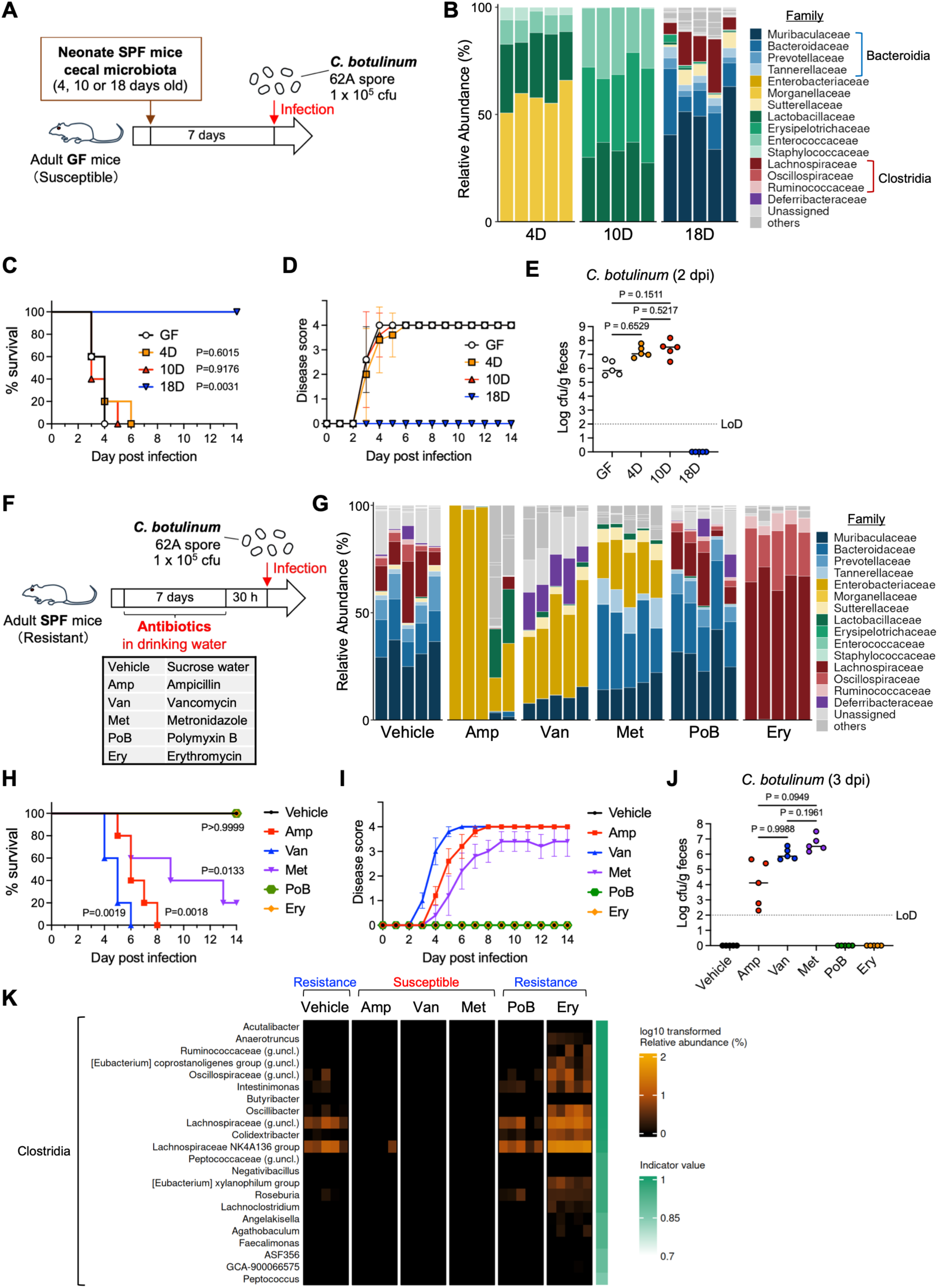
Clostridia are enriched in *C. botulinum*-resistant microbiota. (**A** and **F**) Experimental design. (**B** and **G**) Fecal microbiota composition analyzed by 16S rRNA gene sequencing. (**C** and **H**) Survival after *C. botulinum* infection. (**D** and **I**) Botulism disease scores. Data are presented as mean ± s.d. (**E** and **J**) Fecal *C. botulinum* levels. Horizontal lines indicate the median. (**K**) Indicator species analysis. Only strong resistant-associated indicator genera with IndVal.g ≥ 0.85 are shown. n = 5 mice per group. Statistical analyses were performed using the log-rank (Mantel–Cox) test or one-way ANOVA followed by Tukey’s multiple-comparison test. Data are representative of two independent experiments. dpi, days post infection; LoD, limit of detection.

To clarify the protective role of Clostridia to *C. botulinum* infection, we transplanted GF mice with chloroform-resistant bacteria prepared from adult SPF mouse cecal contents (mCRB), which established a microbiota composed predominantly of Clostridia (Fig. 2A, B) (*12*, *22*). For comparison, we generated gnotobiotic mice colonized with a cocktail of four *Bacteroides* species, another dominant bacterial group in adult mice, consisting of *B. acidifaciens, B. fragilis, B. thetaiotaomicron* and *B. uniformis.* Following *C. botulinum* infection, GF mice and *Bacteroides-*colonized mice developed severe botulism symptoms (Fig. 2C, D). In contrast, mCRB-transplanted mice exhibited complete resistance to *C. botulinum* infection, with no detectable botulism symptoms following challenge. Notably, *C. botulinum* colonization was completely abolished in mCRB-transplanted mice, whereas *Bacteroides*-transplanted mice showed even higher intestinal colonization levels than GF mice (Fig. 2E). CRB derived from adult human feces (hCRB) similarly conferred resistance to *C. botulinum*, suppressing both disease development and intestinal colonization, suggesting a conserved protective mechanism among mammals (Fig. 2B, F–H). Principal coordinate analysis (PCoA) based on unweighted UniFrac distances revealed that microbiota associated with susceptibility clustered separately from resistant microbiota states across the different experimental models (fig. S2). Notably, erythromycin-treated and mCRB-transplanted mice formed a distinct resistant cluster, consistent with the enrichment of Clostridia observed in these groups (Fig. 1G, 2B). Together, these findings demonstrate that commensal Clostridia consortia play a critical role in CR against *C. botulinum*.

**Fig. 2.**
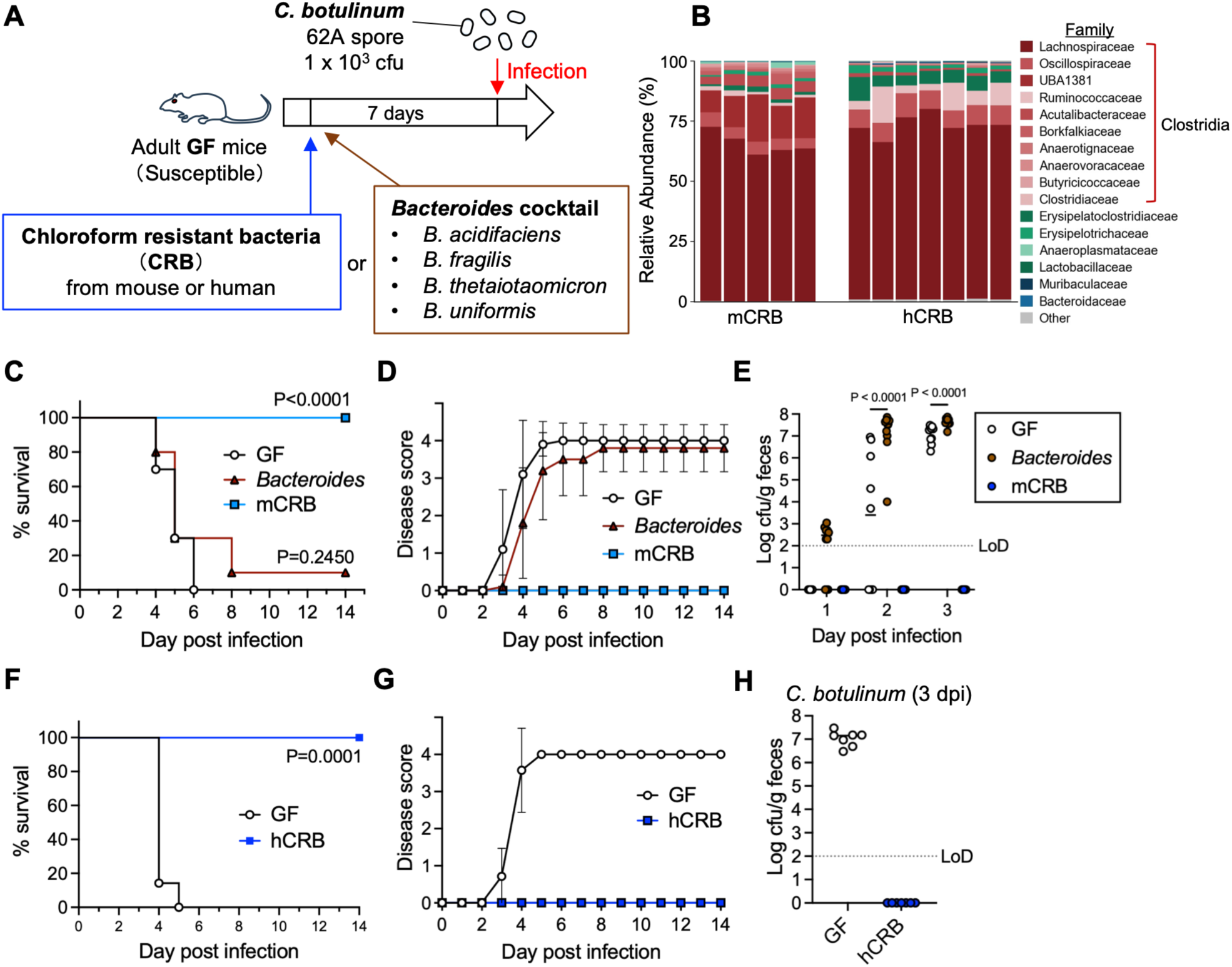
Commensal Clostridia consortia confer resistance to *C. botulinum*. (**A**) Experimental design. (**B**) Fecal microbiota composition analyzed by shotgun metagenomics. (**C** and **F**) Survival after *C. botulinum* infection. (**D** and **G**) Botulism disease scores. Data are presented as mean ± s.d. (**E** and **H**) Fecal *C. botulinum* levels. Horizontal lines indicate the median. n = 10 (**C** to **E**) or 7 (**F** to **H**) mice per group. Statistical analyses were performed using the log-rank (Mantel–Cox) test or one-way ANOVA followed by Tukey’s multiple-comparison test. Data are pooled from two independent experiments. dpi, days post infection; LoD, limit of detection.

### Stickland-metabolizing Clostridia are associated with acquisition of resistance to *C. botulinum*

Although resistance was associated with adult-associated commensal Clostridia, the responsible bacterial species and mechanisms underlying acquisition of resistance during human infant gut microbiota development remained unknown. To address this question, we reconstituted the development of human gut microbiota in adult GF mice by sequentially transplanting fecal samples collected from the same healthy infants over time (Fig. 3A). We termed this experimental model sequential infant fecal microbiota transplantation (siFMT). Although children under 1 year of age are epidemiologically considered susceptible to infant botulism, siFMT mice exhibited a clear transition from susceptibility to complete resistance during microbiota development. In all four infants examined (Infant 1–4), fecal samples collected at earlier time points remained susceptible to *C. botulinum*, permitting intestinal colonization and disease development following infection (Fig. 3B–I, fig. S3A–D). In contrast, samples obtained at later ages conferred complete resistance, with no detectable colonization or botulism symptoms. Notably, resistance emerged abruptly rather than progressively in each individual (Fig. 3F–I), suggesting that acquisition of CR against *C. botulinum* depends on colonization by specific bacteria. To identify bacteria associated with resistance acquisition, we performed shotgun metagenomic analysis of siFMT mice. However, substantial interindividual variation in microbiota composition among infants was observed (fig. S3E–H), making it difficult to identify the responsible bacteria through statistical analysis alone. Consistent with this, neither abundance- nor membership-based PCoA clearly separated resistant and susceptible microbiota (fig. S3I, J). In contrast, metabolomic analysis of feces from siFMT mice identified multiple metabolites that differed between the susceptible and resistant periods, among which the proline-derived bacterial metabolite 5-aminovalerate (5-AVA) showed the most prominent increase during the resistant period (Fig. 3J, K). In all four infants, 5-AVA was scarcely detected during the susceptible period but increased markedly following acquisition of resistance (Fig. 3L–O). Conversely, proline levels decreased during the resistant period, consistent with microbial consumption of proline accompanying increased 5-AVA production (Fig. 3K). These findings identify 5-AVA as a hallmark metabolite of the resistant microbiota and suggest that colonization by 5-AVA-producing bacteria contributes to acquisition of CR against *C. botulinum* in human infants.

**Fig. 3.**
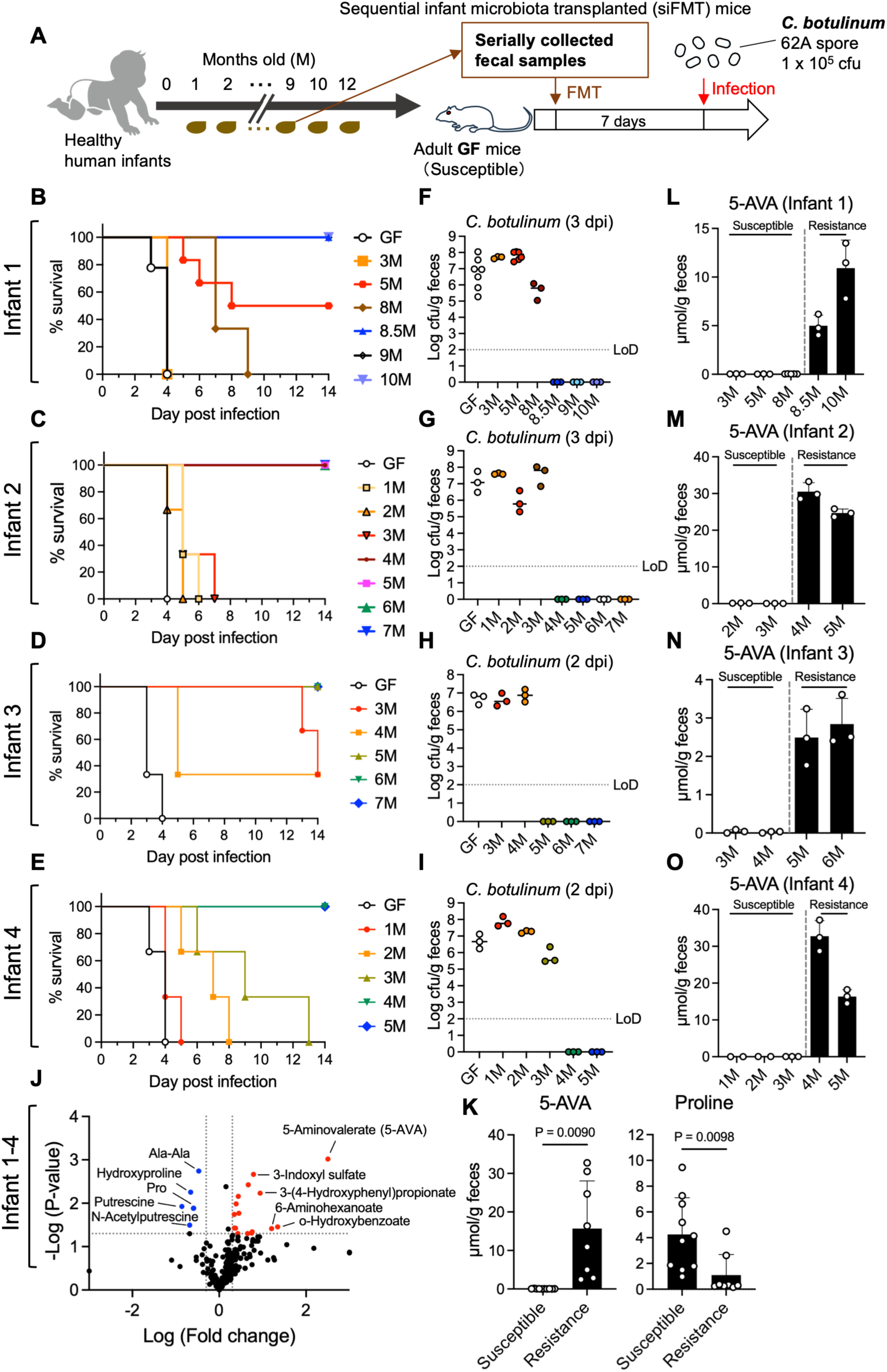
Sequential infant microbiota transplantation reveals acquisition of resistance. (**A**) Experimental design. (**B** to **E**) Survival after *C. botulinum* infection in siFMT mice generated from four individual infants. (**F** to **I**), Fecal *C. botulinum* levels. Each data point represents an individual mouse. Horizontal lines indicate the median. (**J**) Volcano plot of fecal metabolomic analysis comparing susceptible and resistant siFMT mice. Fold changes exceeding the plotting range were clipped to the axis limits. (**K**) Fecal 5-aminovalerate (5-AVA) and proline levels corresponding to the metabolites highlighted in (**J**). Each data point represents the mean value of recipient mice for each infant sampling age. (**L** to **O**) Longitudinal changes in fecal 5-AVA levels in siFMT mice generated from each infant. Each data point represents an individual mouse. For (**K** to **O**), bars indicate mean ± s.d. For infection experiments, n = 3–7 mice per group. For metabolomic comparisons in (**J** and **K**), n = 10 susceptible and 8 resistant infant-age groups. For longitudinal 5-AVA analysis in (**L** to **O**), n = 2–5 mice per group. Statistical analyses were performed using Welch’s t-test. Data are pooled from two or three independent experiments. dpi, days post infection; LoD, limit of detection.

5-AVA is a product of Stickland metabolism, an amino acid metabolic pathway used by a subset of Clostridia to generate energy anaerobically through redox reactions between paired amino acids (*23*, *24*). In this pathway, D-proline is converted to 5-AVA by D-proline reductase (Fig. 4A) (*23*). Consistent with the increased abundance of 5-AVA during the resistant period, metagenomic analysis of feces from siFMT mice revealed enrichment of D-proline reductase (*prd*) genes in resistant microbiota (Fig. 4B, C, fig. S4A). Multiple complementary analyses further supported an association between reductive Stickland metabolism and resistance acquisition (fig. S4B, C). Although most detected *prd* genes were unclassified at the species level, classified sequences identified *Clostridioides difficile*, *Sellimonas intestinalis,* and sequences assigned to *Hungatella hathewayi* AF19-21 as candidate sources of 5-AVA production during the resistant period (Fig. 4D, table S2). We subsequently isolated bacterial strains from resistant fecal samples that showed high 16S rRNA gene sequence similarity to *C. difficile*, *S. intestinalis* and *H. hathewayi* (Fig. 4E). Whole-genome sequencing and ribosomal multilocus sequence typing (rMLST) identified the isolates as non-toxigenic *C. difficile*, *S. intestinalis* and *H. hominis* (Fig. 4E, fig. S5). The *H. hominis* isolate obtained here lacked detectable *prd* genes (table S3). These isolates were individually inoculated into mice transplanted with susceptible infant fecal microbiota (iFMT mice) to examine their effects on susceptibility to *C. botulinum* infection (Fig. 4F). Among them, *C. difficile* conferred complete resistance to *C. botulinum*, preventing both intestinal colonization and disease development (Fig. 4G–I). In contrast, *S. intestinalis* delayed disease onset but failed to prevent *C. botulinum* colonization, and infected mice eventually succumbed to botulism. *H. hominis* showed no detectable protective effect against either colonization or disease development. A commercially available non-toxigenic *C. difficile* strain similarly provided complete protection against both *C. botulinum* colonization and disease development (fig. S6A–D), suggesting that this protective effect represents a species-level trait rather than a strain-specific property. Notably, mono-association with *C. difficile* alone significantly attenuated disease severity but did not prevent *C. botulinum* colonization, indicating that interaction with accessory bacteria is required for complete resistance mediated by *C. difficile* (fig. S6E–H). Collectively, our metabolo-genomic analyses of siFMT mice enabled identification of bacteria conferring resistance to *C. botulinum*.

**Fig. 4.**
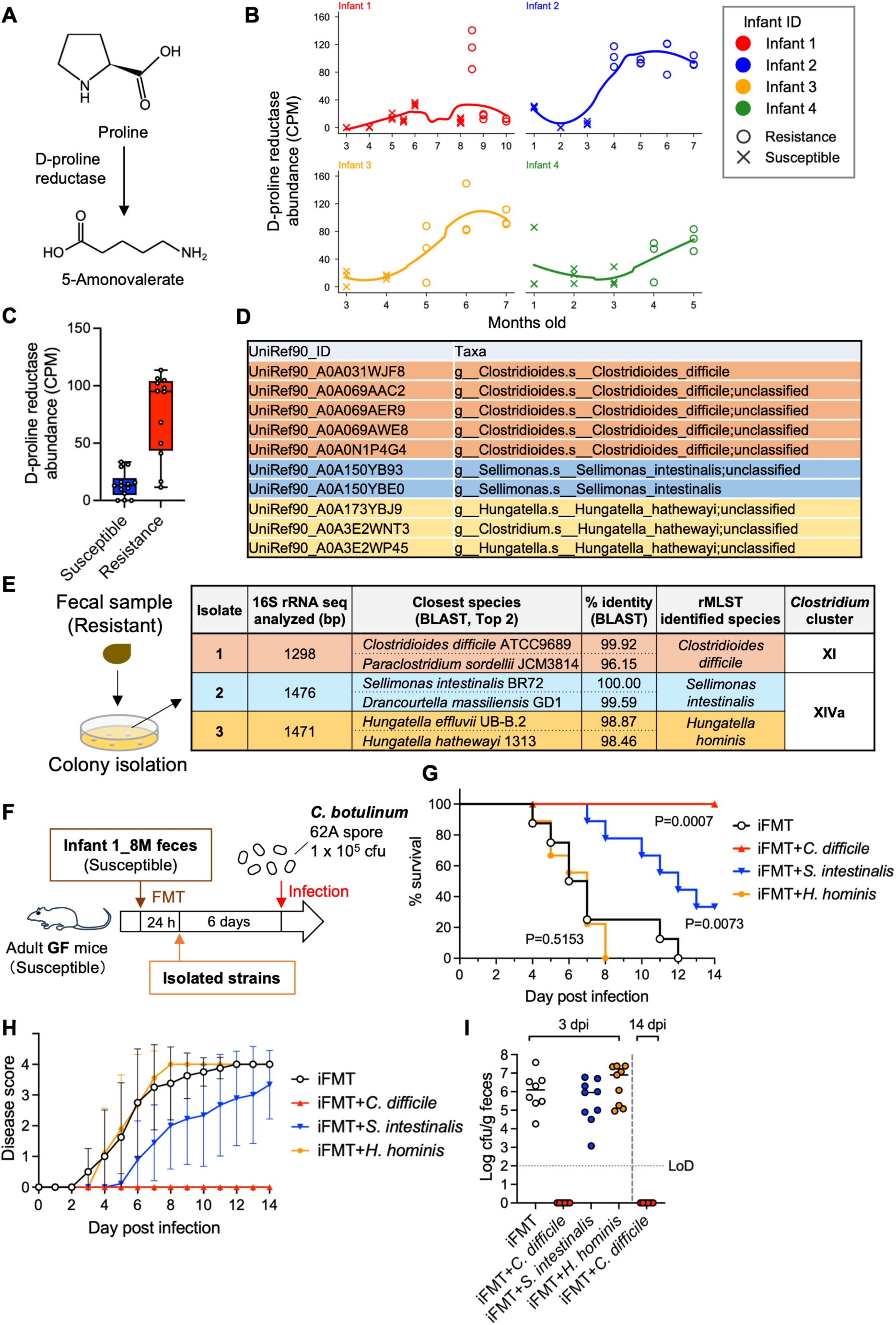
Stickland-metabolizing Clostridia protect against *C. botulinum*. (**A**) Conversion of proline to 5-aminovalerate by D-proline reductase. (**B**) Longitudinal changes in total D-proline reductase gene abundance in siFMT mice shown in Fig. 3. Each data point represents an individual mouse. (**C**) D-proline reductase gene abundance in susceptible and resistant microbiota. Box-and-whisker plots show the median, interquartile range, minimum and maximum values; all data points are shown. Each data point represents the mean value of recipient mice for each infant sampling age. (**D**) Taxonomic assignment of D-proline reductase gene-associated UniRef90 IDs. Representative taxa with species-level annotation are shown. (**E**) Isolation and characterization of bacterial strains from resistant fecal samples. (**F**) Experimental design. (**G**) Survival after *C. botulinum* infection. (**H**) Botulism disease scores. Data are presented as mean ± s.d. (**I**) Fecal *C. botulinum* levels. Each data point represents an individual mouse. Horizontal lines indicate the median. (**F** to **I**) n = 6–9 mice per group. Statistical analyses were performed using the log-rank (Mantel–Cox) test. Data are pooled from three independent experiments. CPM, copies per million; dpi, days post infection; LoD, limit of detection.

### Nutritional niche competition restricts *C. botulinum* colonization

Given that *C. botulinum* itself also possesses Stickland metabolism (*23*), we next hypothesized that competition for substrate amino acids with other Stickland-metabolizing Clostridia underlies CR against *C. botulinum*. To define the metabolic profile of *C. botulinum*, we mono-associated GF mice with a genetically modified non-toxic *C. botulinum* strain (Δ*bontA*), enabling long-term colonization without development of botulism (Fig. 5A). Metabolomic analysis of cecal contents revealed marked alterations in amino acid metabolites following *C. botulinum* Δ*bontA* colonization, indicating extensive amino acid consumption during intestinal colonization (Fig. 5B). Amino acids reduced by colonization included proline, glycine and branched-chain amino acids, whereas 5-AVA, a metabolite associated with Stickland metabolism, was markedly enriched (Fig. 5C, D). We next examined metabolic changes associated with *C. difficile* colonization in the context of infant microbiota. Metabolomic analysis of *C. difficile*-colonized iFMT mice revealed marked enrichment of 5-AVA together with reduced levels of multiple amino acids compared with control iFMT mice (Fig. 5E). Amino acids specifically depleted in *C. difficile*-colonized iFMT mice compared with *S. intestinalis*-or *H. hominis*-colonized iFMT mice included several amino acids associated with Stickland fermentation and vegetative growth (fig. S7A, B), raising the possibility that *C. difficile*-mediated amino acid consumption may restrict expansion of *C. botulinum* within amino-acid-dependent nutritional niches. To directly assess metabolic overlap between these bacteria, we examined the growth of *C. botulinum* in culture supernatants from *C. difficile*. *C. botulinum* growth was impaired in *C. difficile* culture supernatant, whereas amino acid supplementation significantly reduced the growth delay (Fig. 5F, G). These findings further support the idea that *C. difficile* and *C. botulinum* utilize overlapping amino acid resources. Given the marked enrichment of 5-AVA in the resistant microbiota, we examined whether it directly inhibits *C. botulinum* growth. 5-AVA supplementation showed no inhibitory effect *in vitro* (fig. S7C), indicating that 5-AVA is a metabolic byproduct rather than a direct mediator of protection. Together, these findings support the idea that *C. difficile* and *C. botulinum* occupy overlapping amino-acid-dependent nutritional niches in the gut.

**Fig. 5.**
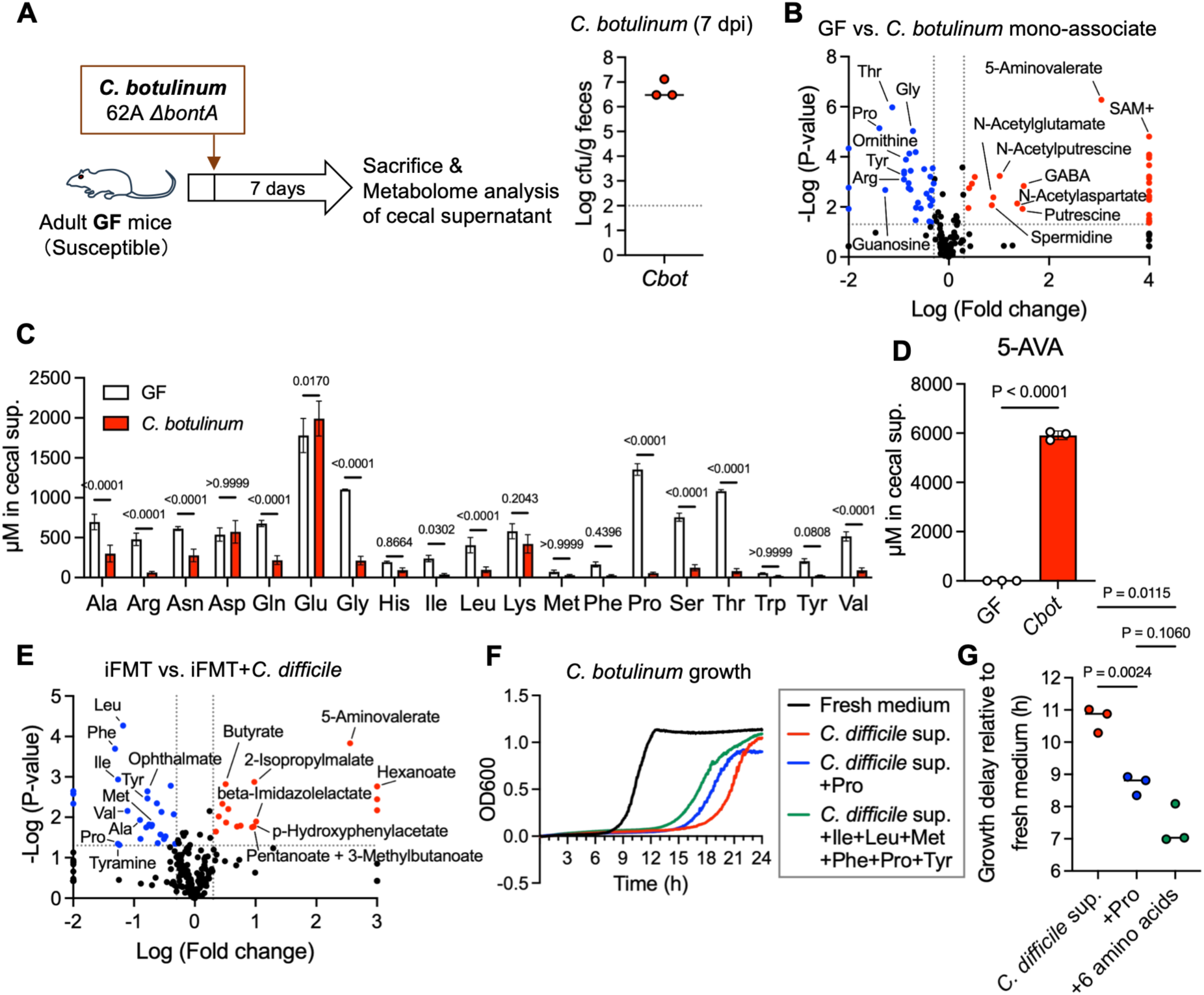
Competition for environmental amino acids underlies resistance to *C. botulinum*. (**A**) Experimental design. (**B**) Volcano plot of cecal metabolomic analysis comparing GF and *C. botulinum* Δ*bontA* mono-associated mice. Fold changes exceeding the plotting range were clipped to the axis limits. (**C** and **D**) Cecal amino acid and 5-aminovalerate (5-AVA) levels in *C. botulinum* Δ*bontA* mono-associated mice. Bars indicate mean ± s.d. (**E**) Volcano plot of fecal metabolomic analysis comparing iFMT mice with or without *C. difficile* colonization shown in Fig. 4. (**F**) Growth of *C. botulinum* in BHIS-S medium or *C. difficile* culture supernatant with or without amino acid supplementation. (**G**) Time required to reach OD600 = 0.5 relative to fresh medium. Each point represents an independent experiment. Horizontal lines indicate the median. n = 3 mice per group (**B** to **E**). Statistical analyses were performed using Welch’s t-test or one-way ANOVA followed by Tukey’s multiple-comparison test. Data in (**F**) are representative of three independent experiments.

Finally, to directly test the importance of amino acid availability for intestinal colonization by *C. botulinum in vivo*, GF mice were fed purified diets containing different concentrations of soy protein as the sole amino acid source (fig. S8A). Mice fed a protein-free diet showed neither detectable *C. botulinum* colonization nor botulism symptoms, whereas mice fed protein-containing diets remained susceptible to infection (fig. S8B–D). Together with the metabolomic and *in vitro* findings, these results support a model in which competition for shared amino-acid-dependent nutritional niches restricts intestinal colonization by *C. botulinum*. Notably, increasing dietary protein content only modestly affected *C. botulinum* colonization levels (fig. S8D), suggesting that amino acid availability in the lower intestine is tightly restricted, thereby creating a nutritional niche in which bacterial competition may occur. Overall, our findings support a model in which acquisition of Stickland-metabolizing Clostridia during infant gut microbiota maturation promotes CR against *C. botulinum* through competition for shared nutritional niches (fig. S9).

## Discussion

The bacterial basis of resistance to infant botulism has remained unresolved since the disease was first recognized nearly five decades ago. Here, we identify acquisition of Stickland-metabolizing Clostridia as a key determinant of resistance to *C. botulinum* colonization. Our findings suggest that occupation of a shared amino-acid-dependent nutritional niche by Stickland-metabolizing Clostridia restricts subsequent *C. botulinum* colonization, providing a mechanistic explanation for how resistance to infant botulism is established during microbiota development. Our study provides two advances with direct relevance to infant botulism: identification of the protective bacteria that establish CR against *C. botulinum*, and discovery of 5-AVA as a candidate biomarker associated with acquisition of CR.

Nutrient competition is one of the most fundamental forms of interaction among bacteria and plays a central role in CR against intestinal pathogens (*25–27*). Our study suggests that resistance to *C. botulinum* is mediated by occupation of a shared amino-acid-dependent nutritional niche by Stickland-metabolizing Clostridia. In these organisms, amino acids serve not only as building blocks for growth but also as substrates for energy generation through Stickland metabolism (*23*, *24*). Because *C. botulinum* relies on this metabolic pathway, it likely competes with established commensal Stickland-metabolizing Clostridia for critical amino-acid resources. Consistent with this model, prior colonization by Stickland-metabolizing Clostridia prevented subsequent *C. botulinum* establishment, supporting a “first-come, first-served” mechanism of CR. Unlike many enteric pathogens that persist at low levels despite CR (*28–31*), *C. botulinum* is completely excluded from resistant microbiota. This unusually strong resistance may reflect the extensive overlap between the metabolic requirements of *C. botulinum* and those of protective commensals. Our dietary manipulation further supported this model, as depletion of dietary amino acids prevented intestinal colonization by *C. botulinum* (fig. S8), consistent with the high dependence of proteolytic *Clostridium* species on environmental amino acids for both growth and energy metabolism (*32–34*). More broadly, our findings suggest that acquisition of specific microbial metabolic functions can shift the gut microbiota from a susceptible to a resistant state by restricting access to shared nutritional niches.

Although *C. difficile* is widely recognized as the major cause of antibiotic-associated diarrhea, approximately 40% of healthy infants and young children are asymptomatic carriers, even when colonized by toxigenic strains (*35*, *36*). Therefore, both toxigenic and non-toxigenic *C. difficile* can be part of the normal gut microbiota in infants; however, the physiological significance of *C. difficile* colonization remains unclear. Recently, a case report showed that colonization by toxigenic *C. difficile* was associated with disappearance of *C. botulinum* in twins with infant botulism (*37*). Because simultaneous detection of *C. difficile* is rare (2.3%) in infant botulism patients, these two species appear unable to stably coexist even in the infant gut (*37*). Consistent with these clinical observations, our results demonstrate that *C. difficile*, previously regarded primarily as a pathogen, contributes to acquisition of resistance against *C. botulinum* in infants. At the same time, complete *C. difficile*-mediated resistance required accessory bacteria in the infant gut microbiota (fig. S6F–H). Because Stickland metabolism can be enhanced by metabolic interactions within microbial communities (*38*), establishment of resistance may depend not only on the presence of Stickland-metabolizing Clostridia but also on community contexts that support their metabolic activity. Importantly, infant microbiota lacking *C. difficile* also acquired resistance to *C. botulinum* (fig. S4A), suggesting that other Stickland-metabolizing Clostridia can also confer resistance. Identifying the bacteria that support *C. difficile* in the resistant microbiota remains an important subject for future investigation. These findings challenge a simple classification of intestinal bacteria as either pathogens or commensals. Although toxigenic Clostridia can cause disease under specific circumstances, our results indicate that *C. difficile* can simultaneously contribute to host protection by excluding more lethal pathogens. Consistent with this notion, non-toxigenic *C. difficile* strains have been explored as live biotherapeutics for prevention of recurrent toxigenic *C. difficile* infection (*39*, *40*). Thus, the ecological functions of gut bacteria may be determined not only by their virulence potential but also by the microbial community context in which they reside.

Several limitations should be considered. First, although siFMT and gnotobiotic mouse models enable causal investigation of microbiota function, they may not fully recapitulate microbial ecology in the human infant intestine. Second, the longitudinal infant cohort was intentionally designed for mechanistic investigation and therefore included a limited number of donors. Third, most experiments were performed using a single group I (proteolytic) *C. botulinum* strain, and the extent to which the identified mechanism applies across the broader diversity of BoNT-producing clostridia remains unclear. Importantly, not all cases of infant botulism are caused by Stickland-metabolizing *C. botulinum*. Other causative species, including *C. butyricum* and *C. baratii*, are primarily saccharolytic organisms that likely depend on ecological mechanisms distinct from those identified here (*41*, *42*). Further studies will therefore be required to elucidate resistance mechanisms against these atypical BoNT-producing bacteria.

In summary, this study reveals a role for nutritional niche competition in microbiota-mediated resistance to *C. botulinum* colonization, establishes a framework for microbiota-based prevention of infant botulism, and highlights the biological significance of microbial metabolic functions acquired during microbiota development.

## Acknowledgements

We thank Dr. Shinichi Nakamura (Kanazawa University) for insightful advice on classical bacteriology. We thank Hitomi Kuraoka, Yuki Konoshita and Sachiyo Akagi (Kanazawa University) for technical assistance. We thank Hiromi Honda (Kanazawa University) for technical assistance and for preparing the schematic illustrations. We thank Dr. Sho Amatsu (Kanazawa University) for establishing a method for genetic modification of *C. botulinum*. We thank Dr. Takumi Nishiuchi (Kanazawa University) for technical support with proteomic analysis. We are grateful to the infants and their parents for participating in this study and providing fecal samples. The authors used ChatGPT (OpenAI) to assist with English language editing, manuscript organization, and improvement of scientific writing. All scientific interpretations, conclusions, and final manuscript content were reviewed and approved by the authors.

## Funding

Japan Agency for Medical Research and Development (AMED) grants JP23fk0108688h0001 and JP25wm0325067h0003 (NK), JP21zf0127001 (SF), and JP19fm0208010h0003 (YF)

Japan Society for the Promotion of Science (JSPS) KAKENHI grants JP25K22552 (NK), JP26H02318 (SF), and JP21H02729 and JP24K02278 (YF)

Kanazawa University “JIKO-CHOKOKU” Project (NK)

Yakult Bio-Science Foundation (NK)

Hokuriku Bank Research Grant for Young Scientists (NK)

The Chemo-Sero-Therapeutic Research Institute (YF)

## Author contributions

Conceptualization: NK, YF

Methodology: NK, YF

Investigation: NK, YK, JY, TM, AY, YA, HT, KI, NH, YN, KS, DM, YM, TW

Formal analysis: NK, YK, JY, DT, HT, SK, DM, YM, SN

Visualization: NK, DT

Funding acquisition: NK, SF, YF

Project administration: NK, YF

Supervision: TM, SF, KH, YF

Writing – original draft: NK

Writing – review & editing: NK, YK, JY, TM, AY, YA, DT, HT, SK, KI, NH, YN, KS, DM, YM, SN, TW, SF, KH, YF

## Competing interests

N.K., Y.F., S.F., J.Y., K.H. and Y.K. are inventors on a pending patent application (Japanese Patent Application No. 2026-112886) related to the findings reported in this study. The remaining authors declare no competing interests.

## Data and materials availability

Sequencing and metabolomic data generated in this study will be deposited in public repositories before publication. Bacterial strains generated in this study are available from the corresponding authors upon reasonable request, subject to institutional biosafety regulations.

## Supplementary Materials

### Materials and Methods

#### Biosafety

*Clostridium botulinum* and botulinum neurotoxins are classified as Class II pathogens under Japanese regulations. All experiments were conducted in accordance with regulations and guidelines approved by the Ministry of Health, Labour and Welfare of Japan and the Kanazawa University Biosafety Committee.

#### Bacterial strains and spore purification

Strains used in this study are listed in table S4. *C. botulinum, C. difficile, S. intestinalis* and *H. hominis* were cultured at 37 °C in an anaerobic chamber (85% nitrogen, 10% hydrogen, 5% carbon dioxide; Concept 400, Baker Ruskinn, Bridgend, UK) in BHIS-S medium, consisting of brain heart infusion (BHI) broth (Shimadzu Diagnostics Corporation, Tokyo, Japan) supplemented with 0.5% (w/v) yeast extract, 4% salts solution, 0.1% L-cysteine, and 0.2% fructose (*43*). The salts solution contained 0.02% CaCl2, 0.02% MgSO4, 0.1% K2HPO4, 0.1% KH2PO4, 1% NaHCO3 and 0.2% NaCl. *B. acidifaciens, B. fragilis, B. thetaiotaomicron* and *B. uniformis* were cultured at 37 °C in an anaerobic chamber in cooked meat medium [0.1 g/ml cooked meat (Kanto Chemical Co., Inc., Tokyo, Japan), 0.3% glucose, and 0.3% soluble starch] for 2 days. *E. coli* was cultured aerobically at 37 °C in LB broth with shaking.

Spores of *C. botulinum* type A strain 62A were purified for use in the intestinal botulism model. Glycerol stocks were inoculated into 3 ml of TPGY broth (5% Tryptone Peptone, 0.5% Peptone, 0.5% Yeast Extract, 0.1% Glucose) and cultured at 37 °C in an anaerobic chamber for 2 days. Cultures were diluted 1:10,000 in sterilized phosphate-buffered saline (PBS) without CaCl2 and MgCl2 (PBS), and 100 µl was spread onto TP agar (5% Tryptone Peptone, 0.5% Peptone, 1.5% agar) plates followed by anaerobic incubation at 37 °C using an AnaeroPack system (Mitsubishi Gas Chemical Company, Inc., Tokyo, Japan) for 8 days. Colonies were scraped from plates with 2 ml of cold sterile water per plate and collected by centrifugation at 8,000 × g for 5 min at 4 °C. Pellets were washed once with 10 ml of cold water and incubated overnight at 4 °C. The washed pellets were resuspended in 5 ml of cold 15% iodixanol (OptiPrep; Serumwerk Bernburg AG, Bernburg, Germany) and gently layered onto 10 ml of 40% iodixanol. Samples were centrifuged at 10,000 × g for 30 min at 4 °C. Pellets were subsequently washed four times with 10 ml of cold water, resuspended in cold water and diluted as appropriate. Purified spores were aliquoted and stored at −80 °C. Before use, spores were heat-treated at 80 °C for 20 min. Colony-forming units (CFU) were determined by plating spores on TPGY agar (TPGY + 1.5% agar) plates and anaerobically culturing at 37 °C for 2 days.

#### Mice

Germ-free (GF) ICR mice were purchased from Japan SLC (Shizuoka, Japan) and maintained in vinyl isolators (JIC Co., Ltd., Tokyo, Japan). Specific pathogen-free (SPF) ICR, BALB/c and *Rag2*^−/−^*Jak3*^−/−^ (BRJ, BALB/c background) mice were purchased from Japan SLC. *Myd88*^−/−^*Trif*^−/−^ mice (C57BL/6 background) were obtained from the Oriental Bio Service (Kyoto, Japan) and maintained under SPF conditions at Kanazawa University. All animal experiments were approved by the Institutional Animal Care and Use Committee of Kanazawa University (AP-214246 and AP-214252) and performed in accordance with institutional guidelines. Female mice aged 6–8 weeks at the time of infection were used as adult mice unless otherwise indicated. For experiments using *Myd88*^−/−^*Trif*^−/−^ mice, both male and female mice were used. Mice were routinely provided with autoclaved water and PicoLab Rodent Diet 20 (PMI Nutrition International, St. Louis, MO, USA). For experiments manipulating intestinal amino acid availability, mice were fed purified diets containing soy protein as the sole amino acid source (Funabashi Farm Co., Ltd., Chiba, Japan) from 1 week before infection until the end of the experiments. The composition of the purified diets is shown in table S5.

#### *C. botulinum* infection in antibiotic-treated mice

A mouse model of intestinal botulism induced by antibiotic treatment was established as previously described (*16*). Briefly, adult SPF mice were administered antibiotics in drinking water for 7 days in individually ventilated cages (IVCs). Ampicillin (Sigma-Aldrich, St. Louis, MO, USA), vancomycin, metronidazole, polymyxin B and erythromycin (FUJIFILM Wako Pure Chemical Corporation, Osaka, Japan) were dissolved individually or as an antibiotic cocktail in sterile drinking water supplemented with 10 g l−1 sucrose. Ampicillin, metronidazole, polymyxin B and erythromycin were used at 1 g l−1, whereas vancomycin was used at 0.5 g l−1. Antibiotic-containing water was sterilized using a 0.22-µm filter and replaced every 3–4 days throughout treatment. Mice were switched to normal drinking water 24–30 h before infection and orally inoculated with 1 × 10^5 CFU of heat-treated *C. botulinum* spores prepared as described above using a feeding needle. Botulism symptoms were assessed daily for 14 days using the following scoring system (*16*): 0, no clinical signs; 1, limb weakness; 2, severe limb weakness; 3, moribund; and 4, dead. Collected fecal samples were stored at −80 °C until use for CFU quantification and bacterial genomic DNA extraction. For quantification of *C. botulinum*, fecal samples were suspended in PBS at 1 mg per 10 µl, serially diluted as appropriate, plated on Brucella HK agar (Kyokuto Pharmaceutical Industrial Co., Ltd., Tokyo, Japan) supplemented with egg yolk, and incubated at 37 °C for 2 days using an AnaeroPack system. Lipase-positive colonies were counted as *C. botulinum* (*16*).

#### Generation and infection of sequential infant fecal microbiota-transplanted (siFMT) mice

Collection of human fecal samples was approved by the Medical Ethics Committee of Kanazawa University (2022–189), and written informed consent was obtained from the parents of all participants. Fecal samples were collected approximately monthly from four healthy Japanese infants of both sexes and anonymized before analysis. Samples were collected in sterile containers, temporarily stored in household freezers, and subsequently stored at −80 °C until processing. Frozen fecal samples were processed in an anaerobic chamber. Samples were diluted 1:40 (w/v) in PBS, passed through a 100-µm cell strainer (Corning Inc., Corning, NY, USA), supplemented with glycerol at a final concentration of 16%, aliquoted, and stored at −80 °C until use. Before transfer into vinyl isolators, aliquots were decontaminated with Acrofine PRO 8000 (JIC). 6-week-old female GF ICR mice were orally inoculated with 200 µl of fecal suspension using a feeding needle. After 1 week of colonization, mice were orally infected with 1 × 10^5 CFU of *C. botulinum* spores as described above and monitored for 14 days. Botulism symptoms and fecal *C. botulinum* levels were evaluated as described for antibiotic-treated mice. For metabolomic analyses, siFMT groups shown in Fig. 3L–O representing microbiota before and after acquisition of resistance to *C. botulinum* were selected based on sample availability.

#### Transplantation of infant mouse microbiota and *C. botulinum* infection

Cecal and colonic contents collected from two 4-day-old SPF ICR mice, or cecal contents collected from individual 10- or 18-day-old SPF ICR mice, were suspended in 5 ml of PBS in an anaerobic chamber and passed through a 100-µm cell strainer. Glycerol was added to a final concentration of 10%, and aliquoted samples were stored at −80 °C until use. Before transfer into vinyl isolators, frozen samples were decontaminated for more than 30 min as described above. 6-week-old female GF ICR mice were orally inoculated with 200 µl of microbiota suspension using a feeding needle. One week after transplantation, mice were orally infected with 1 × 10^5 CFU of *C. botulinum* spores as described above. Botulism symptoms and fecal *C. botulinum* levels were evaluated as described for antibiotic-treated mice.

#### Preparation and transplantation of chloroform-resistant bacteria (CRB)

CRB were prepared from human and mouse microbiota as previously described with slight modifications (*12*, *22*). Whole intestinal tracts were collected from a 7-week-old female SPF ICR mouse and immediately transferred into an anaerobic chamber. One fecal pellet from the upper colon and an equal amount of cecal contents were suspended in 18 ml of anaerobic PBS and vigorously vortexed. For human CRB, fecal samples collected from a healthy 34-year-old human adult male were diluted 1:40 (w/v) in PBS and passed through a 100-µm cell strainer before chloroform treatment. 14 ml of each suspension was transferred into a Hungate tube, and chloroform was added to a final concentration of 3%. Tubes were sealed with butyl rubber stoppers, removed from the anaerobic chamber, vigorously shaken by hand for 5 min, and incubated at 37 °C with rotation for 1 h. After incubation, residual chloroform was removed by bubbling CO2 gas through the suspension for 10 min using a sterile gas line equipped with a 0.2-µm filter and needles inserted through the butyl rubber stopper. The resulting suspension was decontaminated with Acrofine PRO 8000 before transfer into vinyl isolators. Adult female GF mice were orally inoculated with 500 (mCRB) or 300 µl (hCRB) of the chloroform-treated suspension using a feeding needle. One week after transplantation, mice were orally infected with 1 × 10^3 CFU of *C. botulinum* spores as described above.

#### Colonization of mice with bacterial cultures and *C. botulinum* infection

Cultures of bacterial isolates or commercially available strains were decontaminated with Acrofine PRO 8000 before transfer into vinyl isolators. For administration of bacterial cocktails, cultures were mixed at equal volumes in an anaerobic chamber immediately before use. Adult GF mice or infant fecal microbiota-transplanted mice were orally inoculated with 500 µl of bacterial culture using a feeding needle. One week after bacterial inoculation, mice were orally infected with 1 × 10^3–10^5 CFU of *C. botulinum* spores as described above. Botulism symptoms and fecal *C. botulinum* levels were evaluated as described for antibiotic-treated mice. For metabolomic analysis of *C. botulinum* Δ*bontA* mono-associated mice, cecal contents were collected 1 week after colonization, suspended in an equal volume of PBS, and centrifuged at 10,000 × g for 2 min. To remove viable pathogens before sample transfer, the supernatants were filtered through 0.22-µm membrane filters and subjected to metabolomic analysis.

#### Isolation of bacteria from resistant fecal samples

For isolation of *C. difficile*, infant fecal samples collected during the resistant period were suspended in BHIS-S medium at 100 mg ml−1, heat-treated at 65 °C for 20 min, and cultured in cooked meat medium at 37 °C for 4 days using an AnaeroPack system. Cultures were serially diluted and plated onto Cycloserine-Cefoxitin Mannitol Agar (CCMA) plates followed by anaerobic incubation at 37 °C for 2 days. Individual colonies were screened by colony PCR using *C. difficile*-specific 23S rRNA gene primers. For isolation of *S. intestinalis* and *H. hominis*, fecal samples from hCRB-transplanted mice were diluted in PBS and plated onto BHIS-S agar plates followed by anaerobic incubation at 37 °C for 2 days in an anaerobic chamber. A total of 196 randomly selected colonies were cultured individually and screened by 16S rRNA gene sequencing. For colony PCR, bacterial colonies or cultures were suspended in 50 mM NaOH and heated at 95 °C for 10 min. After addition of one-tenth volume of 1 M Tris-HCl (pH 8.0), lysates were used as templates for PCR using KOD One PCR Master Mix (TOYOBO Co., Ltd., Osaka, Japan). PCR products were analyzed by Sanger sequencing (Eurofins Genomics K.K., Tokyo, Japan), and bacterial identities were assigned by BLAST analysis. Isolates corresponding to *C. difficile*, *S. intestinali*s, and *H. hominis* were further purified by two rounds of single-colony isolation and stored as glycerol stocks at −80 °C. The resulting isolates were designated KZ3001, KZ3002, and KZ3003, respectively. Specific primers are listed in table S6.

#### Whole-genome sequencing, assembly and phylogenetic analysis

Whole-genome sequencing was performed using a combination of short-read sequencing on the NovaSeq X Plus platform (Illumina, Inc., San Diego, CA, USA) and long-read sequencing on the MinION platform (Oxford Nanopore Technologies, Oxford, UK). For short-read sequencing, DNA libraries were prepared using the Illumina DNA PCR-Free Prep kit according to the manufacturer’s instructions and sequenced in 151-bp paired-end mode on the NovaSeq X Plus platform. For long-read sequencing, 1 μg of genomic DNA was sheared using a g-TUBE device (Covaris, LLC, Woburn, MA, USA) targeting approximately 8-kb fragments. Sequencing libraries were prepared using the Native Barcoding Kit (SQK-NBD114; Oxford Nanopore Technologies) and sequenced on the MinION platform using an FLO-MIN114 flow cell. Long reads were assembled de novo using Flye version 2.7. The assembled contigs were subsequently polished with Illumina short reads using BWA version 0.7.17 for read alignment and Pilon version 1.23 for error correction. Whole-genome phylogenetic analysis was performed using the Type (Strain) Genome Server (TYGS), an automated platform for genome-based taxonomy and phylogenomic inference (*44*). Pairwise genome comparisons were conducted using the Genome BLAST Distance Phylogeny (GBDP) approach, and intergenomic distances were calculated using the trimming algorithm and distance formula d5. A whole-genome phylogenetic tree was inferred from the resulting distances using FastME version 2.1.6.1 with subtree pruning and regrafting (SPR) postprocessing. The tree was midpoint-rooted and visualized using PhyD3.

#### Generation of the *bontA*-deficient *C. botulinum*

A *bontA*-deficient mutant of *C. botulinum* 62A was generated by homologous recombination as previously described (*45*). Briefly, approximately 1.5-kb regions upstream and downstream of *bontA* were amplified from genomic DNA of *C. botulinum* 62A and fused by splicing by overlap extension PCR (SOE-PCR). The resulting fragment was cloned into the BamHI/PstI sites of pXMTL. The plasmid was transferred into *C. botulinum* by conjugation using *E. coli* CA434 as the donor strain. Single- and double-crossover recombinants were selected by antibiotic selection and xylose counterselection. Deletion of *bontA* was confirmed by colony PCR and Sanger sequencing. Mutant strains were stored in TPGY broth containing 18% (w/v) glycerol at −80 °C. Specific primers are listed in table S6.

#### In vitro growth assay of C. botulinum

A glycerol stock of the *C. difficile* isolate was inoculated into 5 ml BHIS-S medium and cultured at 37 °C for 2 days in an anaerobic chamber. The culture was centrifuged at 8,000 × g for 3 min, and the resulting supernatant was filter-sterilized before use. Growth of *C. botulinum* was examined in BHIS-S medium or *C. difficile* culture supernatant supplemented with PBS or amino acids. For amino acid supplementation, isoleucine, leucine, methionine, phenylalanine, proline and tyrosine were added to final concentrations of 1.5, 4, 0.5, 2.5, 0.6 and 0.8 mM, respectively, based on previously reported amino acid uptake by *C. difficile*(*46*) and amino acids specifically depleted in *C. difficile*-colonized infant fecal microbiota-transplanted (iFMT) mice in our metabolomic analysis (fig. S7A). Each condition was dispensed into 96-well plates at 200 µl per well in duplicate. *C. botulinum* 62A spores were heat-treated at 80 °C for 20 min, cooled on ice and added to each well at 1 × 10^4 CFU per well. Plates were incubated statically at 37 °C in an anaerobic chamber, and bacterial growth was monitored by measuring optical density (OD600) over time using a microplate reader Stratus (Cerillo, Charlottesville, VA, USA). To evaluate the inhibitory effect of 5-aminovalerate, growth assays were similarly performed in BHI medium supplemented with PBS or 50 mM 5-aminovalerate (final concentration; FUJIFILM Wako Pure Chemical).

#### 16S rRNA gene amplicon sequencing and analysis

Approximately 50 mg of mouse feces was transferred to a 2-ml tube containing 0.1-mm and 3.0-mm zirconia beads. After addition of InhibitEX buffer from the QIAamp Fast DNA Stool Mini Kit (QIAGEN, Hilden, Germany), samples were mechanically disrupted using a Shake Master Neo (1,500 rpm, 10 min; Bio Medical Science, Tokyo, Japan), and genomic DNA was extracted according to the manufacturer’s protocol. The V3–V4 region of the bacterial 16S rRNA gene was amplified using primers 341F (5′-CCTACGGGNGGCWGCAG-3′) and 805R (5′-GACTACHVGGGTATCTAATCC-3′), followed by paired-end sequencing on the Illumina MiSeq platform (2 × 300 bp). Raw sequence data were processed using the nf-core/ampliseq pipeline (v2.16.1). Primer sequences were removed with Cutadapt (v5.2), and amplicon sequence variants (ASVs) were inferred using DADA2 (v1.34.0) (*47*). Taxonomic assignment was performed against the SILVA database (release 138.2) (*48*). A phylogenetic tree was constructed in QIIME 2 (v2024.10) (*49*) using MAFFT alignment and FastTree 2. Downstream analyses were performed in R (v4.4.2) using the phyloseq and vegan packages. Family-level taxonomic composition was calculated from relative abundances and visualized as stacked bar plots. Beta diversity was assessed using unweighted UniFrac distances (*50*) and visualized by principal coordinate analysis (PCoA). Differences in community composition were evaluated by permutational multivariate analysis of variance (PERMANOVA) (*51*) with 999 permutations. Indicator taxa associated with resistance were identified using the indicspecies package with the group-equalized indicator value statistic (IndVal.g) (*52*) and 999 permutations.

#### Shotgun metagenomic analysis

DNA was extracted from approximately 50 mg of mouse feces using the QIAamp PowerFecal Pro DNA Kit (QIAGEN, Hilden, Germany) with mechanical disruption using a Shake Master Neo (1,500 rpm, 10 min). Shotgun metagenomic libraries were sequenced on an Illumina NovaSeq platform (2 × 150 bp paired-end). After removal of host-derived reads by alignment to the mouse reference genome (GRCm39), taxonomic profiling was performed using Kraken2 (*53*) and Bracken (*54*) with the Mouse Gastrointestinal Bacterial Catalogue (MGBC) database (*55*). Marker-gene-based taxonomic profiles were additionally generated using MetaPhlAn 4 (*56*). Community composition analyses were performed in R using the phyloseq and vegan packages. Relative abundances were visualized as stacked bar plots. Beta diversity was assessed using Bray–Curtis and Jaccard distances and visualized by principal coordinate analysis (PCoA). Functional profiling was performed using HUMAnN 3 (*57*) with the UniRef90 database. Genes involved in Stickland metabolism were quantified from normalized gene-family abundances.

#### Metabolomic analysis

Metabolites in cecal and fecal samples were extracted as previously described (*58*). Briefly, samples were lyophilized using a freeze dryer (TAITEC Corporation, Saitama, Japan) and disrupted with a dispensing spoon. Ten-milligram samples were combined with four 3.0-mm zirconia/silica beads and 100 mg of 0.1-mm beads (TOMY SEIKO Co., Ltd., Tokyo, Japan), and metabolites were extracted in 500 µl methanol by vigorous shaking at 1,500 rpm for 5 min using a Shake Master NEO. Samples were further mixed with 200 µl Milli-Q water and 500 µl chloroform containing 20 µM methionine sulfone, 20 µM 2-(N-morpholino)ethanesulfonic acid (MES), and 20 µM D-camphor-10-sulfonic acid (CSA) as internal standards for capillary electrophoresis time-of-flight mass spectrometry (CE–TOFMS).

Samples were centrifuged at 4,600×g for 30 min at 4 °C. Supernatants were transferred to a 5-kDa-cutoff filter column (Ultrafree MC-PLHCC 250/pk for Metabolome Analysis, Human Metabolome Technologies, Inc., Tsuruoka, Japan) and centrifuged at 9,100×g overnight at 4 °C. Filtrates were evaporated using a CentriVap Centrifugal Vacuum Concentrator (Labconco Corporation, Kansas City, MO, USA). Dried samples were stored at −80 °C and subsequently dissolved in 50 µl Milli-Q water containing 200 µM trimesate and 3-aminopyrrolidine before analysis. CE–TOFMS analyses were performed in both positive and negative ion modes as previously described (*59–61*). An Agilent capillary electrophoresis system (Agilent Technologies, Santa Clara, CA, USA) was used for all CE–TOFMS analyses. Detected peak areas were normalized to methionine sulfone for cationic metabolites and CSA for anionic metabolites. Peak detection and metabolite quantification were performed using MasterHands version 2.19.0.1.63.

#### Proteomic analysis

Plasma and cecal contents collected from two mice per group were subjected to proteomic analysis. Mice were anesthetized and blood was collected by cardiac puncture into heparinized tubes. Plasma was isolated by centrifugation and diluted in PBS containing cOmplete protease inhibitor cocktail (Roche, Basel, Switzerland). Cecal contents were suspended in PBS containing protease inhibitor cocktail, centrifuged, and sequentially filtered through 0.45-µm and 0.22-µm membrane filters. To inactivate botulinum neurotoxin (BoNT), samples were heated at 95 °C for 10 min before storage at −80 °C. Protein samples were dried, resuspended in 6 M urea and 50 mM triethylammonium bicarbonate, reduced with tris(2-carboxyethyl)phosphine, alkylated with iodoacetamide, and digested with trypsin (Promega Corporation, Madison, WI, USA). Peptides were desalted using StageTips and analyzed using an EASY-nLC 1200 system coupled to an Orbitrap Q Exactive Plus mass spectrometer (Thermo Fisher Scientific, Waltham, MA, USA). Peptides were separated on an Aurora C18 column (IonOpticks, Melbourne, Australia) using a linear acetonitrile gradient in 0.1% formic acid. MS/MS spectra were analyzed using the SEQUEST HT algorithm implemented in Proteome Discoverer v2.2 (Thermo Fisher Scientific) against the *Mus musculus* UniProt database. Oxidation of methionine was set as a variable modification and carbamidomethylation of cysteine as a fixed modification. False discovery rates were estimated using a target–decoy strategy and filtered at 1%. Label-free quantification was performed using precursor ion intensities normalized by total peptide amount. Proteins showing >2-fold increases were included in Gene ontology (GO) analyses irrespective of statistical significance because of the exploratory nature of the study. GO enrichment analyses were performed using ShinyGO v0.85.1. Upregulated proteins are listed in table S7.

#### Statistics

Statistical analyses were performed using GraphPad Prism (GraphPad Software, Boston, MA, USA) and Microsoft Excel (Microsoft Corporation, Redmond, WA, USA). Differences between two groups or among multiple groups were analyzed using Welch’s t-test or one-way ANOVA followed by Tukey’s multiple-comparison test, respectively. Survival curves were analyzed using the Kaplan–Meier method with the log-rank (Mantel–Cox) test.

---

**Fig. S1.**
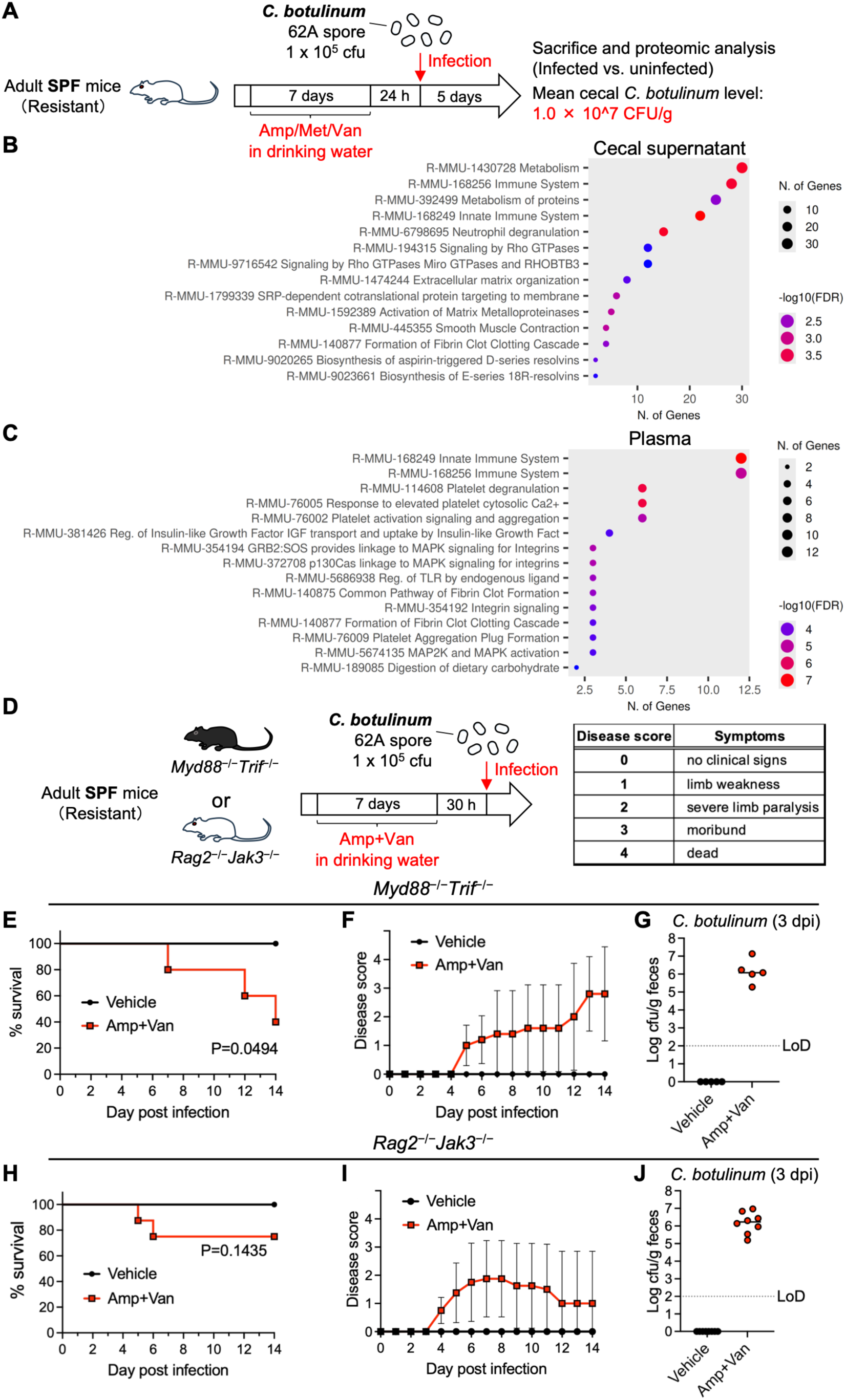
Colonization resistance protects against *C. botulinum* independently of host immunity. (**A** and **D**) Experimental design. (**B** and **C**) Gene ontology analysis of proteomic features elevated in cecal supernatants (**B**) or plasma (**C**) from *C. botulinum*-infected mice compared with uninfected controls. (**E** and **H**) Survival after *C. botulinum* infection. (**F** and **I**) Botulism disease scores. Data are presented as mean ± s.d. (**G** and **J**) Fecal *C. botulinum* levels. Horizontal lines indicate the median. n = 5 (**E** to **G**) or 8 (**H** to **J**) mice per group. Statistical analyses were performed using the log-rank (Mantel–Cox) test. Data are representative (**E** to **G**) or pooled (**H** to **J**) from two independent experiments. dpi, days post infection; LoD, limit of detection.

**Fig. S2.**
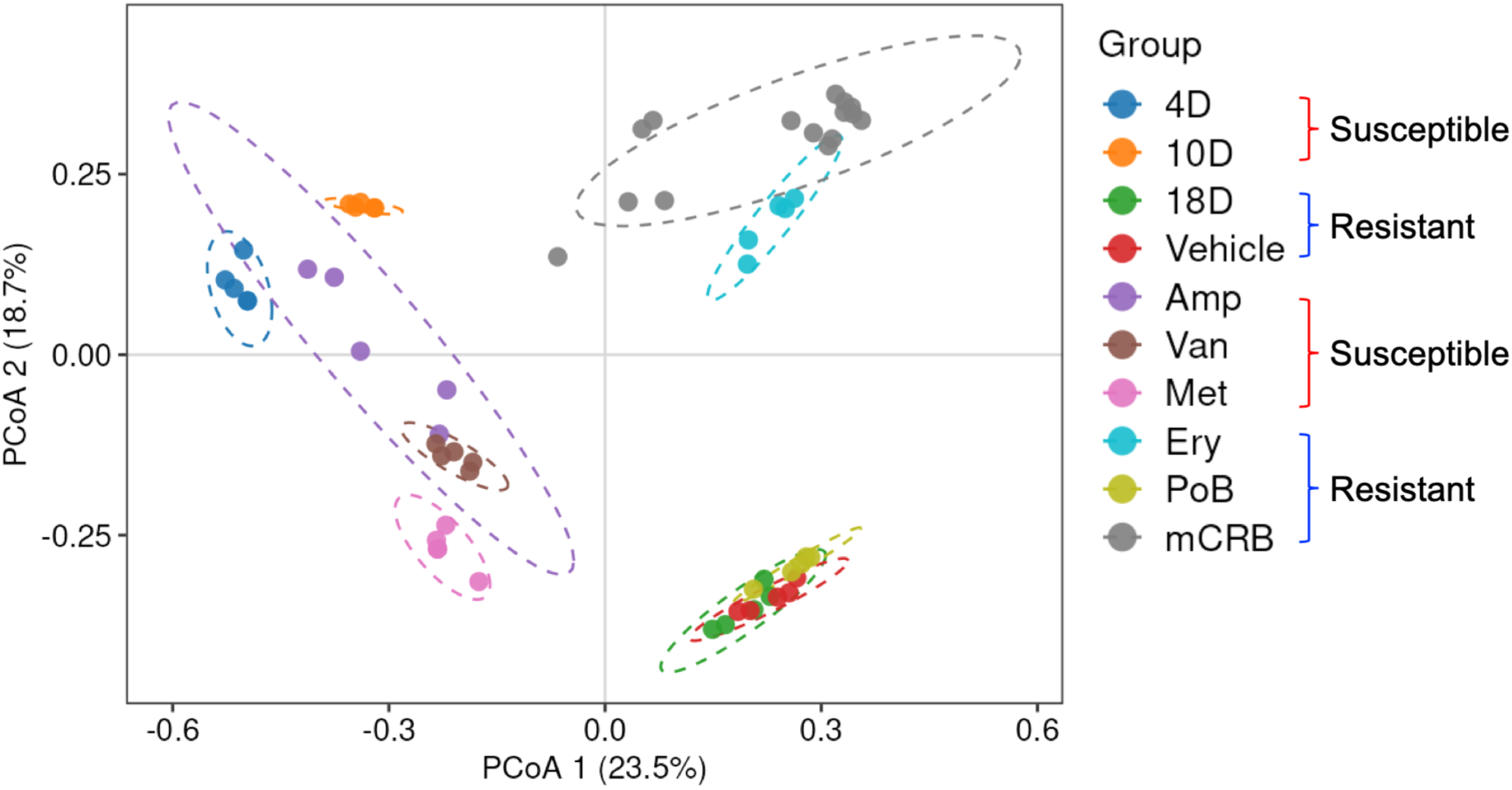
Distinct microbiota structures associated with susceptible and resistant states. Principal coordinate analysis (PCoA) based on unweighted UniFrac distances of 16S rRNA gene amplicon sequencing data from the experiments shown in Fig. 1 and Fig. 2. Each point represents one mouse fecal sample. Colors indicate treatment groups. Ellipses denote 95% confidence intervals.

**Fig. S3.**
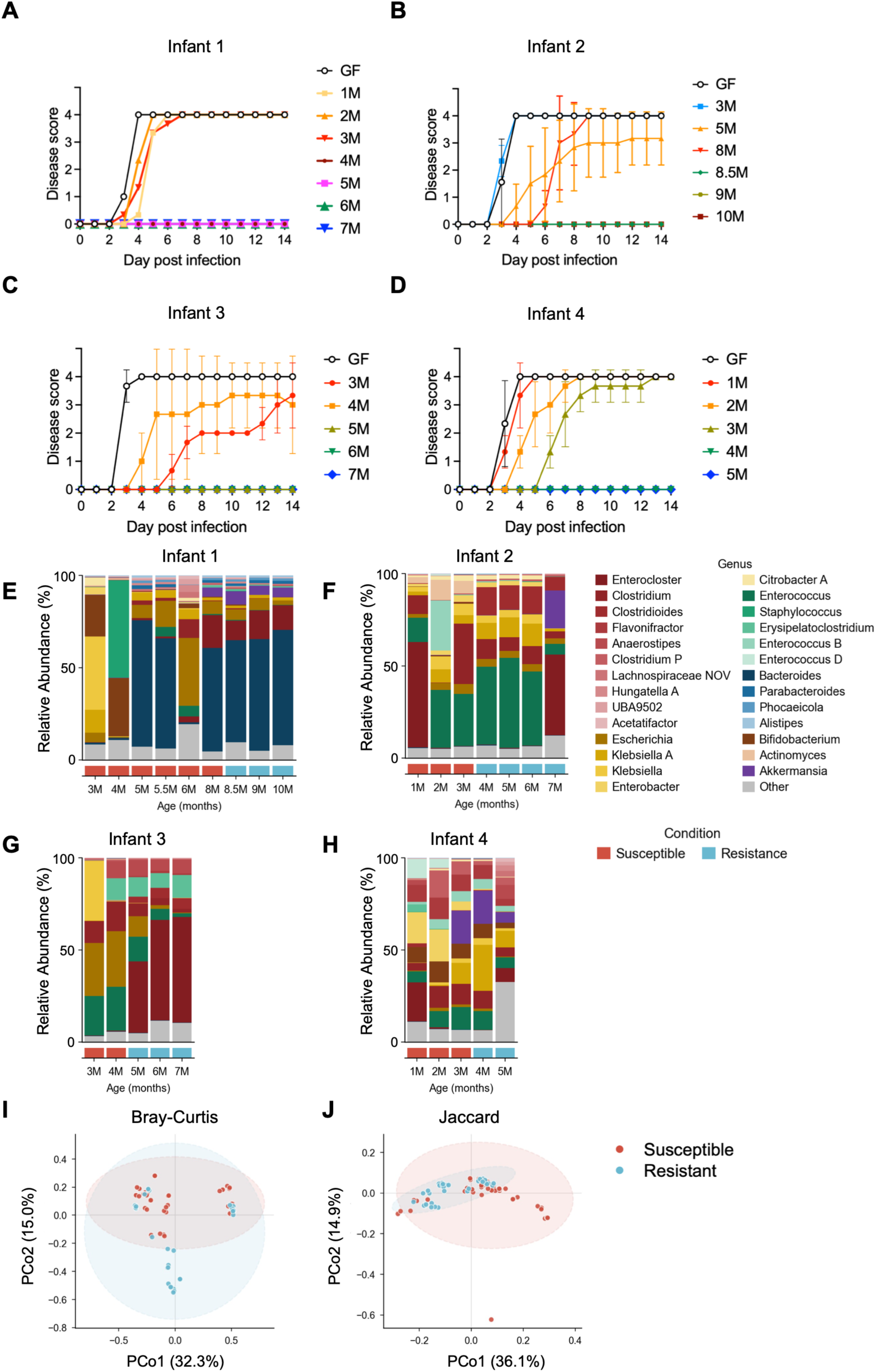
Disease phenotypes and microbiota composition in siFMT mice. (**A** to **D**) Botulism disease scores of siFMT mice shown in Fig. 3. Data are presented as mean ± s.d. (**E** to **H**) Fecal microbiota composition analyzed by shotgun metagenomics. Bars represent the mean relative abundance of siFMT mice at each sampling age from individual infants. (**I** and **J**) Principal coordinate analysis (PCoA) of shotgun metagenomic profiles using Bray–Curtis dissimilarity (**I**) and Jaccard distance (**J**). Each point represents one siFMT mouse. Colors indicate resistant and susceptible groups. Ellipses denote 95% confidence intervals.

**Fig. S4.**
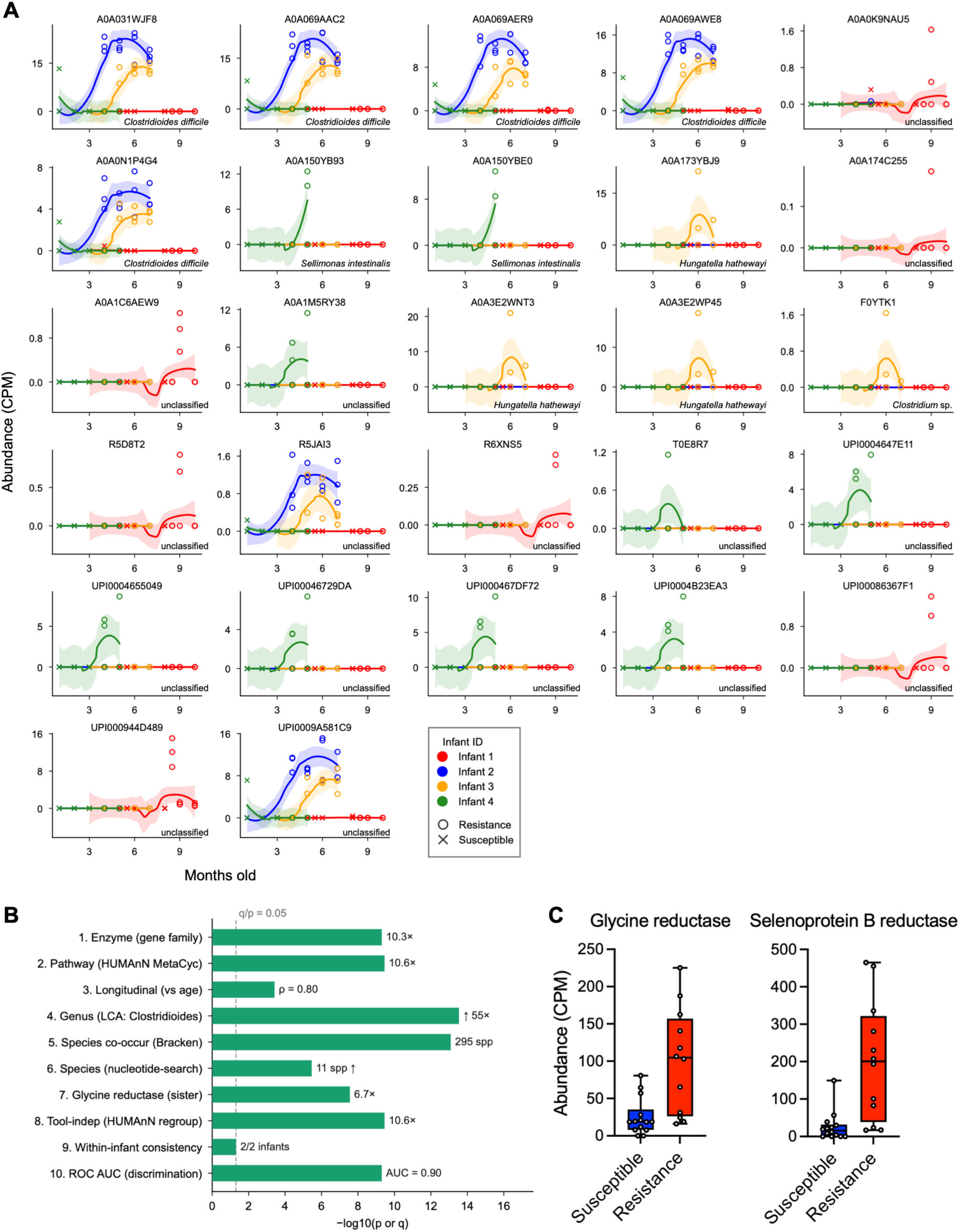
Reductive Stickland metabolism is consistently associated with resistance acquisition. (**A**) Longitudinal changes in abundance of individual D-proline reductase gene-associated UniRef90 IDs contributing to the total D-proline reductase gene abundance shown in Fig. 4B. Each data point represents an individual mouse. (**B**) Ten complementary analyses performed at the gene, pathway and taxonomic levels consistently identified reductive Stickland metabolism as a feature associated with resistance acquisition. Values represent the significance or effect size obtained using each analytical approach. (**C**) Glycine reductase and selenoprotein B reductase gene abundance in susceptible and resistant microbiota. Box-and-whisker plots show the median, interquartile range, minimum and maximum values; all data points are shown. Each data point represents the mean value of recipient mice for each infant sampling age. CPM, copies per million.

**Fig. S5.**
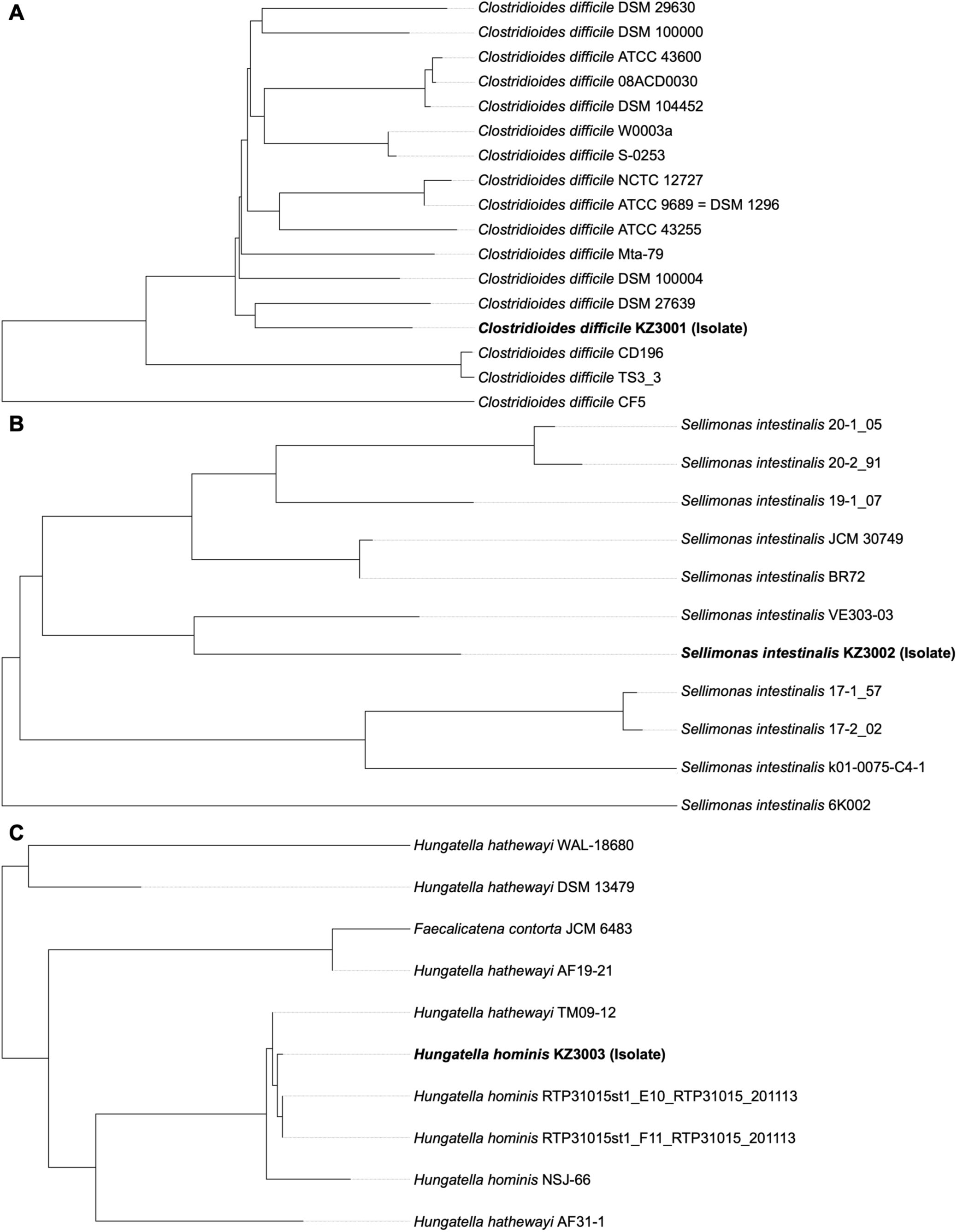
Phylogenetic placement of the isolates. Whole-genome phylogenetic tree inferred using FastME version 2.1.6.1 from Genome BLAST Distance Phylogeny (GBDP) distances calculated from genome sequences in TYGS. Branch lengths are scaled according to GBDP distance formula d5. The tree was rooted at the midpoint. Isolates obtained in this study (KZ3001–KZ3003) are indicated in bold. (**A**) *C. difficile* KZ3001. (**B**) *S. intestinali*s KZ3002. (**C**) *H. hominis* KZ3003.

**Fig. S6.**
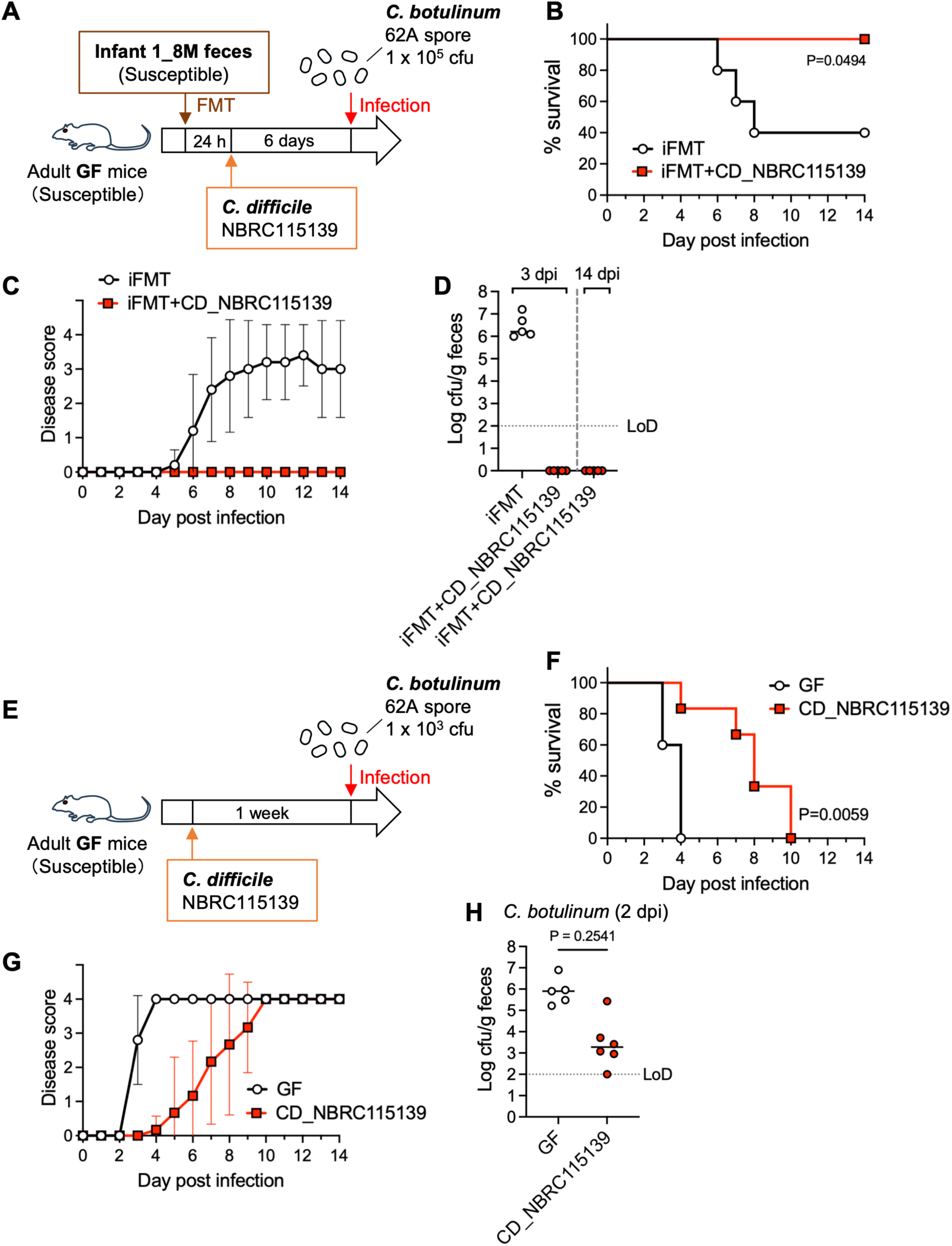
A commercially available *C. difficile* strain confers resistance to *C. botulinum*. (**A** and **E**) Experimental design. (**B** and **F**) Survival after *C. botulinum* infection. (**C** and **G**) Botulism disease scores. Data are presented as mean ± s.d. (**D** and **H**) Fecal *C. botulinum* levels. Horizontal lines indicate the median. n = 5 (**B** to **D**) or 5–6 (**F** to **H**) mice per group. Statistical analyses were performed using the log-rank (Mantel–Cox) test. Data are representative (**B** to **D**) or pooled (**F** to **H**) from two independent experiments. dpi, days post infection; LoD, limit of detection.

**Fig. S7.**
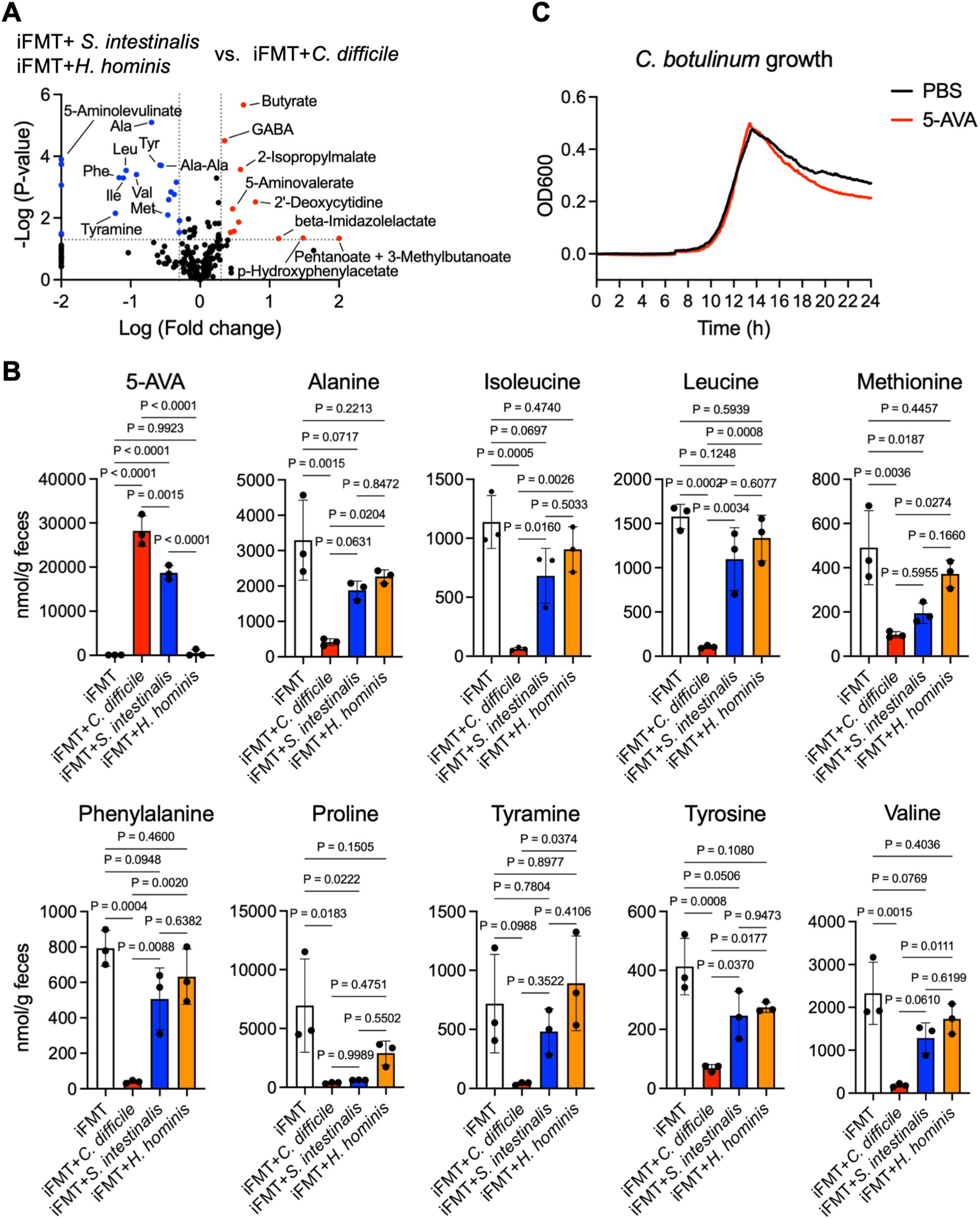
Metabolic signatures associated with *C. difficile* colonization. (**A**) Volcano plot of fecal metabolomic analysis comparing *C. difficile*-colonized iFMT mice with susceptible iFMT mice colonized with *S. intestinalis* or *H. hominis* shown in Fig. 4. Fold changes exceeding the plotting range were clipped to the axis limits. (**B**) Fecal 5-aminovalerate (5-AVA) and amino acid levels in isolate-colonized iFMT mice. Amino acids significantly reduced in *C. difficile*-colonized mice in (**A**) are shown. Bars indicate mean ± s.d. (**C**) Growth of *C. botulinum* in BHI medium with or without 5-AVA supplementation. n = 3 mice per group (**A** and **B**). Statistical analyses were performed using Welch’s t-test or one-way ANOVA followed by Tukey’s multiple-comparison test. Data in (**C**) are representative of two independent experiments.

**Fig. S8.**
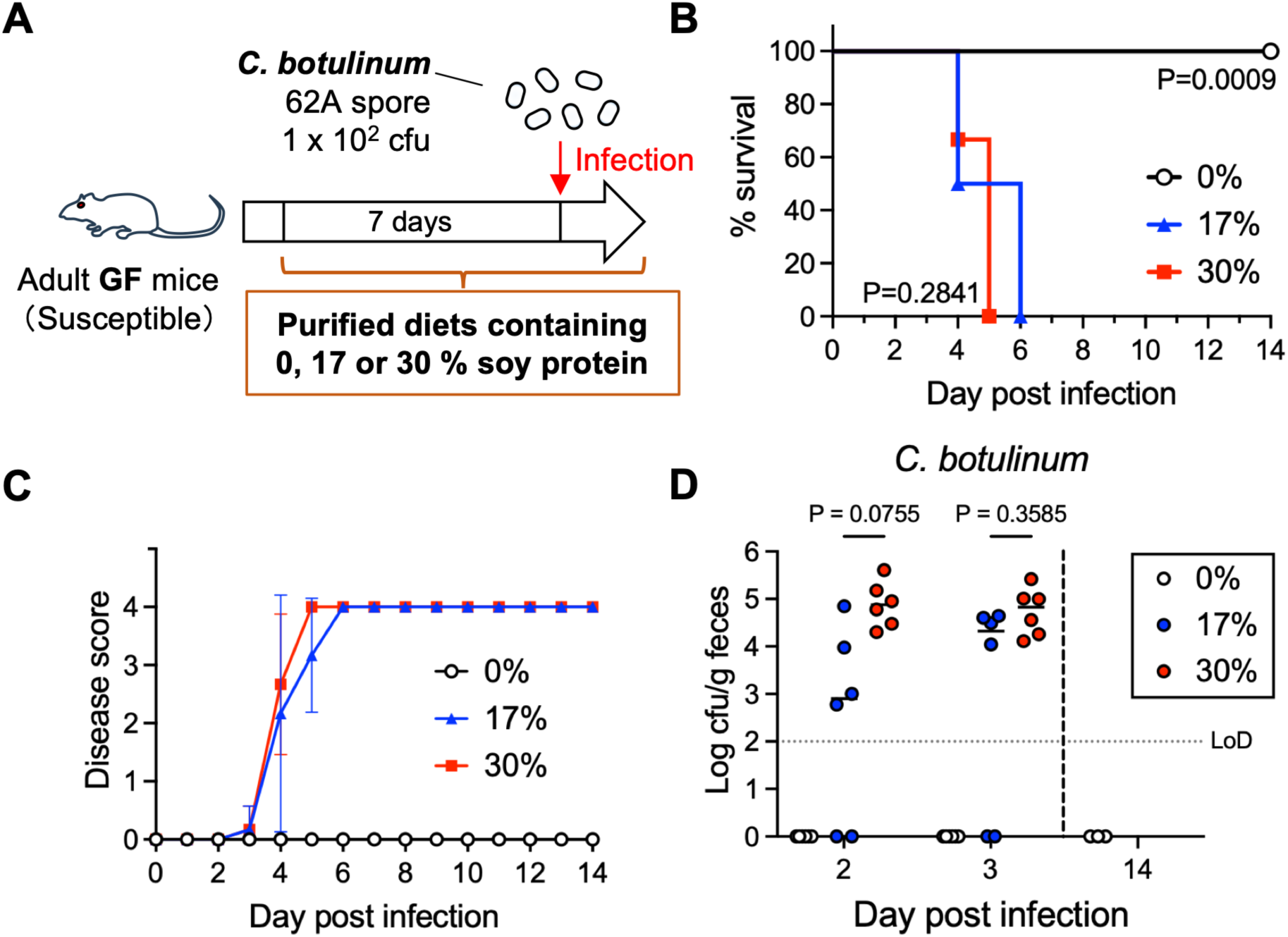
Dietary amino acid availability determines intestinal colonization by *C. botulinum*. (**A**) Experimental design. (**B**) Survival after *C. botulinum* infection. (**C**) Botulism disease scores. Data are presented as mean ± s.d. (**D**) Fecal *C. botulinum* levels. Horizontal lines indicate the median. n = 6 mice per group. Statistical analyses were performed using the log-rank (Mantel–Cox) test or one-way ANOVA followed by Tukey’s multiple-comparison test. Data are pooled from two independent experiments. LoD, limit of detection.

**Fig. S9.**
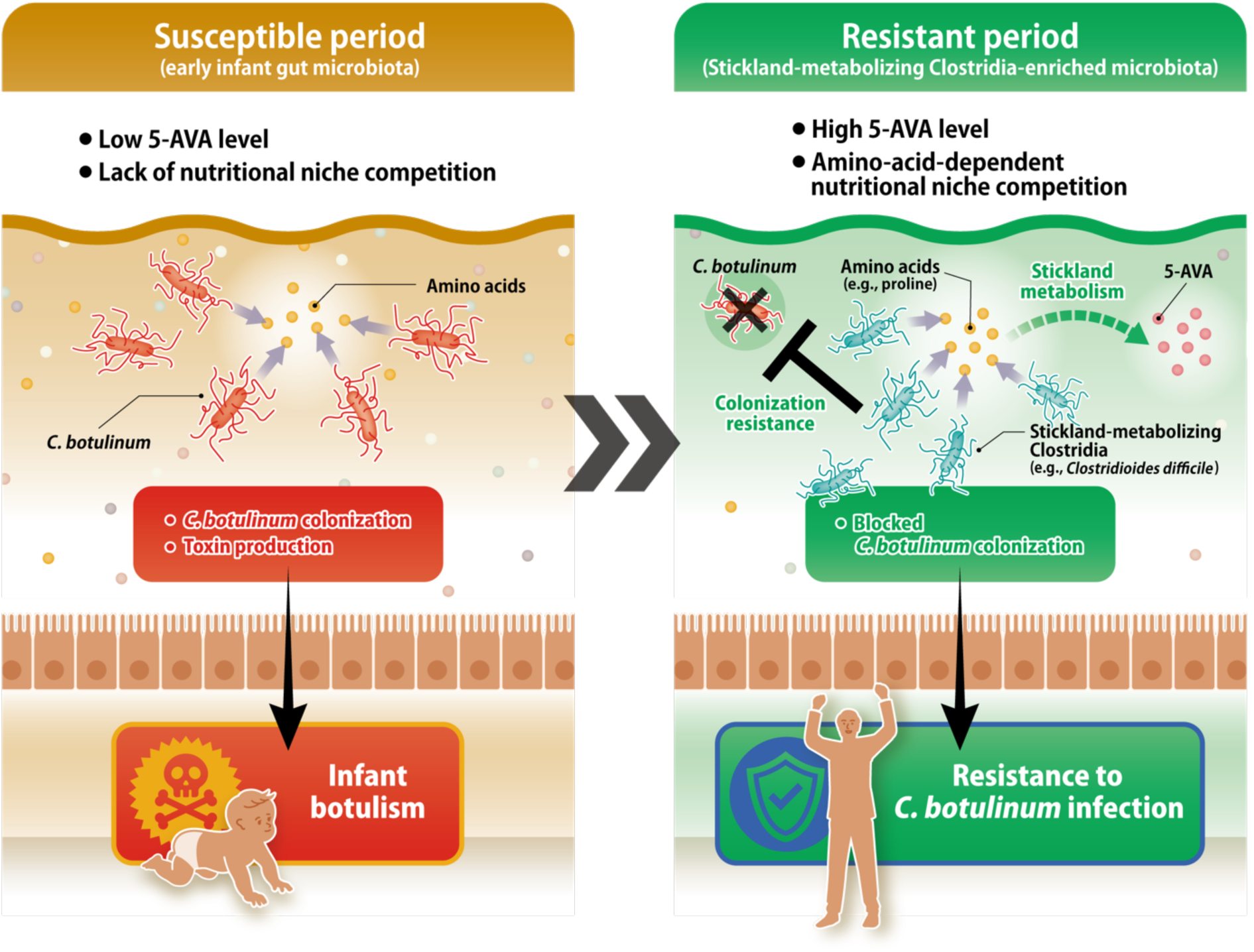
Model of resistance acquisition during infant gut microbiota maturation. Schematic model illustrating microbiota-dependent acquisition of resistance to *C. botulinum* during infant gut microbiota development. During the susceptible period, *C. botulinum* gains access to amino-acid-dependent nutritional niches in the lower intestine, permitting bacterial expansion and toxin production, resulting in infant botulism. During the resistant period, acquisition of Stickland-metabolizing Clostridia establishes amino-acid-dependent nutritional niche competition, thereby restricting access of *C. botulinum* to shared nutritional niches and preventing infection. Increased 5-aminovalerate (5-AVA) levels reflect Stickland metabolic activity associated with the resistant microbiota.

**Table S1.** Indicator taxa associated with the resistant microbiota identified by indicator species analysis.

| Genus | Family | Class | Phylum | IndVal.g | p.value |
| --- | --- | --- | --- | --- | --- |
| Acutalibacter | Ruminococcaceae | Clostridia | Bacillota | 1 | 0.001 |
| Anaerotruncus | Ruminococcaceae | Clostridia | Bacillota | 1 | 0.001 |
| Ruminococcaceae (g.uncl.) | Ruminococcaceae | Clostridia | Bacillota | 1 | 0.001 |
| [Eubacterium] coprostanoligenes group (g.uncl.) | [Eubacterium] coprostanoligenes group | Clostridia | Bacillota | 1 | 0.001 |
| Oscillospiraceae (g.uncl.) | Oscillospiraceae | Clostridia | Bacillota | 1 | 0.001 |
| Intestinimonas | Oscillospiraceae | Clostridia | Bacillota | 1 | 0.001 |
| Butyribacter | Lachnospiraceae | Clostridia | Bacillota | 1 | 0.001 |
| Oscillibacter | Oscillospiraceae | Clostridia | Bacillota | 0.99966997 | 0.001 |
| Lachnospiraceae (g.uncl.) | Lachnospiraceae | Clostridia | Bacillota | 0.9984681 | 0.001 |
| Colidextribacter | Oscillospiraceae | Clostridia | Bacillota | 0.99581599 | 0.001 |
| Lachnospiraceae NK4A136 group | Lachnospiraceae | Clostridia | Bacillota | 0.99201925 | 0.001 |
| Peptococcaceae (g.uncl.) | Peptococcaceae | Clostridia | Bacillota | 0.96609178 | 0.001 |
| Negativibacillus | Ruminococcaceae | Clostridia | Bacillota | 0.96609178 | 0.001 |
| [Eubacterium] xylanophilum group | Lachnospiraceae | Clostridia | Bacillota | 0.96609178 | 0.001 |
| Roseburia | Lachnospiraceae | Clostridia | Bacillota | 0.96609178 | 0.001 |
| Lachnoclostridium | Lachnospiraceae | Clostridia | Bacillota | 0.96609178 | 0.001 |
| Angelakisella | Ruminococcaceae | Clostridia | Bacillota | 0.93094934 | 0.001 |
| Agathobaculum | Butyricicoccaceae | Clostridia | Bacillota | 0.93094934 | 0.001 |
| Faecalimonas | Lachnospiraceae | Clostridia | Bacillota | 0.93094934 | 0.001 |
| ASF356 | Lachnospiraceae | Clostridia | Bacillota | 0.89442719 | 0.001 |
| GCA-900066575 | Lachnospiraceae | Clostridia | Bacillota | 0.89442719 | 0.001 |
| Peptococcus | Peptococcaceae | Clostridia | Bacillota | 0.85634884 | 0.001 |
| Alistipes | Rikenellaceae | Bacteroidia | Bacteroidota | 0.81649658 | 0.001 |
| UCG-010 (g.uncl.) | UCG-010 | Clostridia | Bacillota | 0.81649658 | 0.001 |
| Lachnospiraceae FCS020 group | Lachnospiraceae | Clostridia | Bacillota | 0.81649658 | 0.001 |
| Marvinbryantia | Lachnospiraceae | Clostridia | Bacillota | 0.81649658 | 0.001 |
| Acetatifactor | Lachnospiraceae | Clostridia | Bacillota | 0.81649658 | 0.001 |
| Prevotellaceae UCG-001 | Prevotellaceae | Bacteroidia | Bacteroidota | 0.81636064 | 0.001 |
| unclassified (g.uncl.) | NA | Clostridia | Bacillota | 0.81420124 | 0.001 |
| unclassified (g.uncl.) | NA | Bacilli | Bacillota | 0.77459667 | 0.001 |
| Anaerovoracaceae (g.uncl.) | Anaerovoracaceae | Clostridia | Bacillota | 0.77459667 | 0.001 |
| Family XIII AD3011 group | Anaerovoracaceae | Clostridia | Bacillota | 0.77459667 | 0.001 |
| Lachnospiraceae UCG-010 | Lachnospiraceae | Clostridia | Bacillota | 0.77459667 | 0.002 |
| Desulfovibrio | Desulfovibrionaceae | Desulfovibrionia | Thermodesulfobacteriota | 0.77099817 | 0.004 |
| A2 | Lachnospiraceae | Clostridia | Bacillota | 0.76899008 | 0.001 |
| Butyricimonas | Marinifilaceae | Bacteroidia | Bacteroidota | 0.73029674 | 0.001 |
| Paludicola | Ruminococcaceae | Clostridia | Bacillota | 0.73029674 | 0.003 |
| [Eubacterium] siraeum group | Ruminococcaceae | Clostridia | Bacillota | 0.73029674 | 0.005 |
| UCG-009 | Butyricicoccaceae | Clostridia | Bacillota | 0.73029674 | 0.001 |
| Lachnospiraceae UCG-006 | Lachnospiraceae | Clostridia | Bacillota | 0.7188741 | 0.017 |
| Anaerotignum | Lachnospiraceae | Clostridia | Bacillota | 0.68313005 | 0.011 |
| Ruthenibacterium | Ruminococcaceae | Clostridia | Bacillota | 0.63245553 | 0.02 |
| NK4A214 group | Oscillospiraceae | Clostridia | Bacillota | 0.57735027 | 0.042 |

**Table S2.**
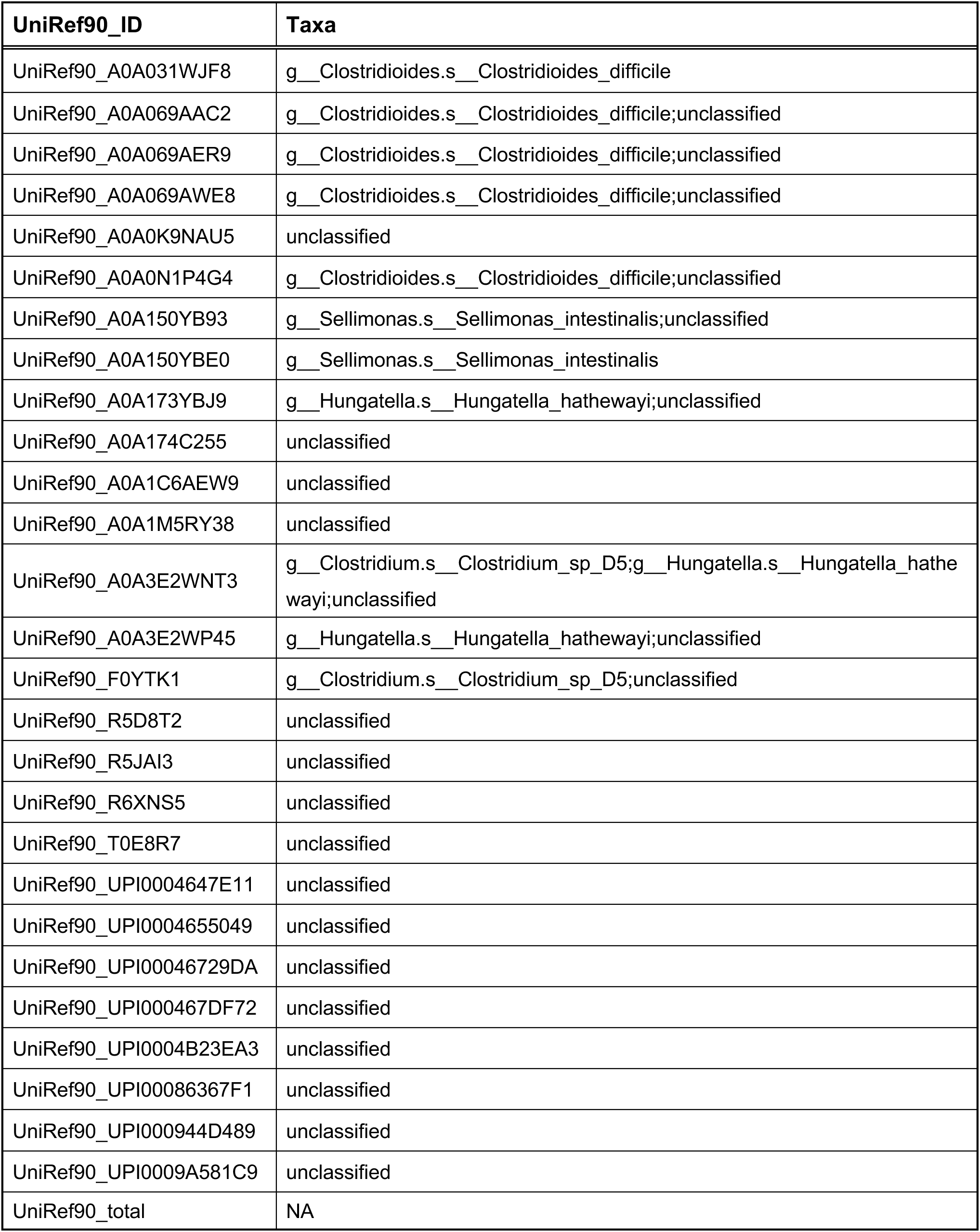
Taxonomic assignment of detected D-proline reductase genes.

| UniRef90_ID | Taxa |
| --- | --- |
| UniRef90_A0A031WJF8 | g__Clostridioides.s__Clostridioides_difficile |
| UniRef90_A0A069AAC2 | g__Clostridioides.s__Clostridioides_difficile;unclassified |
| UniRef90_A0A069AER9 | g__Clostridioides.s__Clostridioides_difficile;unclassified |
| UniRef90_A0A069AWE8 | g__Clostridioides.s__Clostridioides_difficile;unclassified |
| UniRef90_A0A0K9NAU5 | unclassified |
| UniRef90_A0A0N1P4G4 | g__Clostridioides.s__Clostridioides_difficile;unclassified |
| UniRef90_A0A150YB93 | g__Sellimonas.s__Sellimonas_intestinalis;unclassified |
| UniRef90_A0A150YBE0 | g__Sellimonas.s__Sellimonas_intestinalis |
| UniRef90_A0A173YBJ9 | g__Hungatella.s__Hungatella_hathewayi;unclassified |
| UniRef90_A0A174C255 | unclassified |
| UniRef90_A0A1C6AEW9 | unclassified |
| UniRef90_A0A1M5RY38 | unclassified |
| UniRef90_A0A3E2WNT3 | g__Clostridium.s__Clostridium_sp_D5;g__Hungatella.s__Hungatella_hathewayi;unclassified |
| UniRef90_A0A3E2WP45 | g__Hungatella.s__Hungatella_hathewayi;unclassified |
| UniRef90_F0YTK1 | g__Clostridium.s__Clostridium_sp_D5;unclassified |
| UniRef90_R5D8T2 | unclassified |
| UniRef90_R5JAI3 | unclassified |
| UniRef90_R6XNS5 | unclassified |
| UniRef90_T0E8R7 | unclassified |
| UniRef90_UPI0004647E11 | unclassified |
| UniRef90_UPI0004655049 | unclassified |
| UniRef90_UPI00046729DA | unclassified |
| UniRef90_UPI000467DF72 | unclassified |
| UniRef90_UPI0004B23EA3 | unclassified |
| UniRef90_UPI00086367F1 | unclassified |
| UniRef90_UPI000944D489 | unclassified |
| UniRef90_UPI0009A581C9 | unclassified |
| UniRef90_total | NA |

**Table S3.**
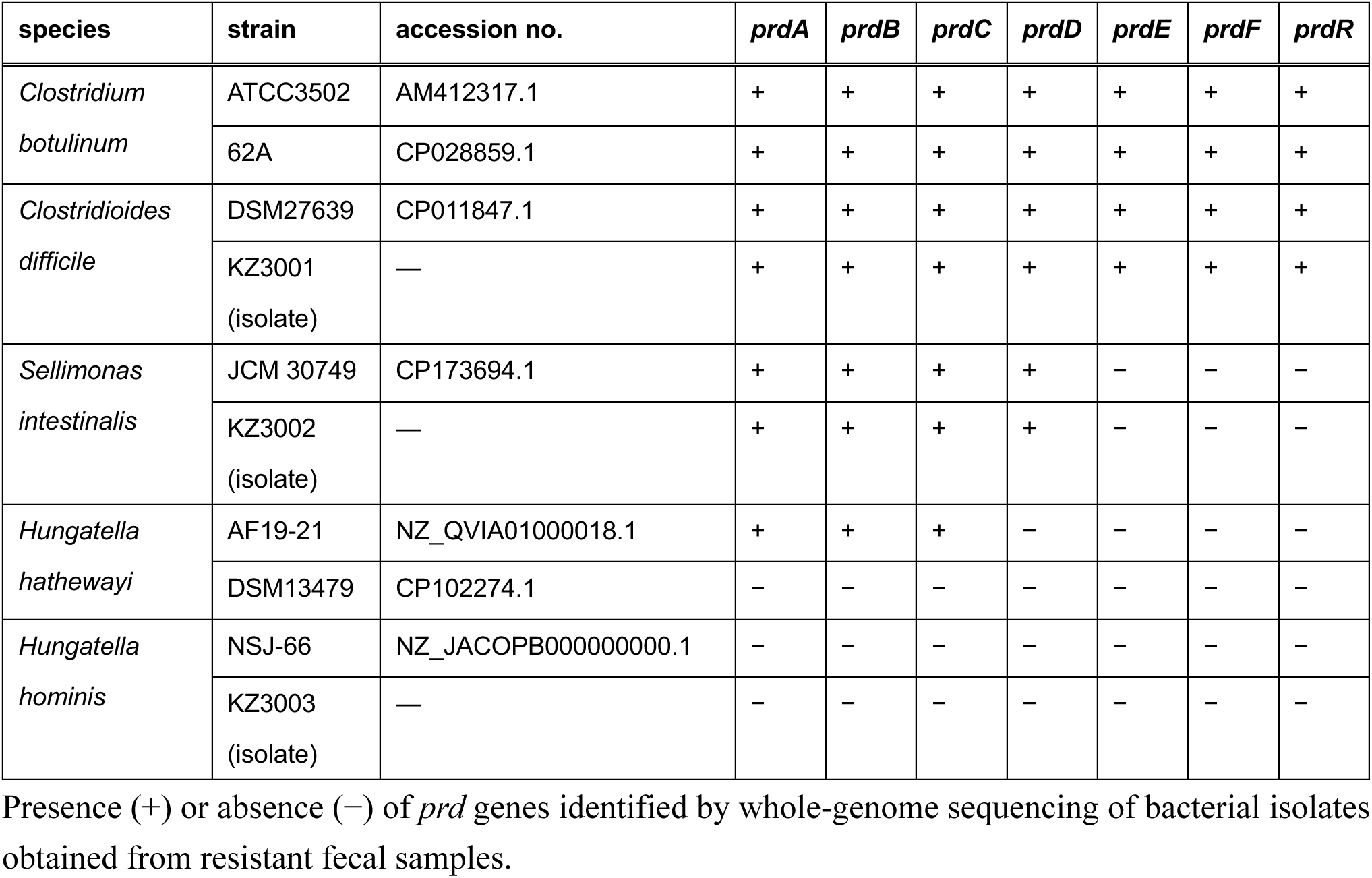
Presence of proline reductase (*prd*) genes in bacterial isolates associated with resistance.

**Table S4.** Bacterial strains used in this study.

| Bacterial strain | Relevant feature or genotype | Source |
| --- | --- | --- |
| *Clostridium botulinum* 62A | Group I, Type A | Laboratory collection |
| *Clostridium botulinum* 62A Δ*bontA* | *bontA* knockout. Constructed as described previously (45). | This study |
| *Bacteroides acidifaciens* JCM10556 | Type strain | JCM |
| *Bacteroides fragilis* JCM11019 | Type strain | JCM |
| *Bacteroides thetaiotaomicron* JCM5827 | Type strain | JCM |
| *Bacteroides uniformis* JCM5828 | Type strain | JCM |
| *Clostridioides difficile* NBRC115139 | TcdA and TcdB negative | NBRC |
| *Clostridioides difficile* KZ3001 | Isolate from resistant fecal sample.<br>TcdA and TcdB negative. | This study |
| *Sellimonas intestinalis* KZ3002 | Isolate from resistant fecal sample | This study |
| *Hungatella hominis* KZ3003 | Isolate from resistant fecal sample | This study |

JCM, Japan Collection of Microorganisms, RIKEN BioResource Research Center, Tsukuba, Japan;
NBRC, Biological Resource Center, National Institute of Technology and Evaluation, Chiba, Japan.

**Table S5.** Composition of purified diets.

**A. Ingredient**
| Ingredient | Unit | 0% soy protein | 17% soy protein | 30% soy protein |
| --- | --- | --- | --- | --- |
| Soy protein | % | — | 17 | 30 |
| β-corn starch | % | 52.15 | 45.05 | 29.8 |
| α-corn starch | % | 9.9 | — | 2.25 |
| Maltodextrin | % | 12.5 | 12.5 | 12.5 |
| Sucrose | % | 10 | 10 | 10 |
| Cellulose powder | % | 5 | 5 | 5 |
| Corn oil | % | 5 | 5 | 5 |
| AIN-76 mineral mixture | % | 3.5 | 3.5 | 3.5 |
| Sodium bicarbonate | % | 0.75 | 0.75 | 0.75 |
| AIN-76A vitamin mixture | % | 1 | 1 | 1 |
| Choline bitartrate | % | 0.2 | 0.2 | 0.2 |
| Total | % | 100 | 100 | 100 |

**B. Nutritional composition**
| Nutritional composition (in 100 g) | Unit | 0% soy protein | 17% soy protein | 30% soy protein |
| --- | --- | --- | --- | --- |
| Moisture | g | 9 | 9 | 9 |
| Crude protein | g | 0.1 | 16.5 | 25.5 |
| Crude fat | g | 5.3 | 5.1 | 5.1 |
| Crude ash | g | 3.3 | 3.2 | 4.3 |
| Crude fiber | g | 4.9 | 4.8 | 4.8 |
| Nitrogen-free extract | g | 77.4 | 61.3 | 51.3 |
| Calories | kcal | 357.5 | 357.4 | 353 |
| Protein calorie ratio | %/kcal | 0.1 | 18.5 | 28.9 |
| Fat calorie ratio | %/kcal | 13.3 | 12.9 | 12.9 |
| Nitrogen-free extract calorie ratio | %/kcal | 86.6 | 68.6 | 58.2 |
| Ca | g | 0.49 | 0.48 | 0.48 |
| P | g | 0.39 | 0.39 | 0.39 |
| Mg | g | 0.05 | 0.05 | 0.05 |
| Na | g | 0.3 | 0.3 | 0.67 |
| K | g | 0.36 | 0.35 | 0.35 |

Purified diets were manufactured by Funabashi Farm and sterilized by 30-kGy gamma irradiation.

**Table S6.**
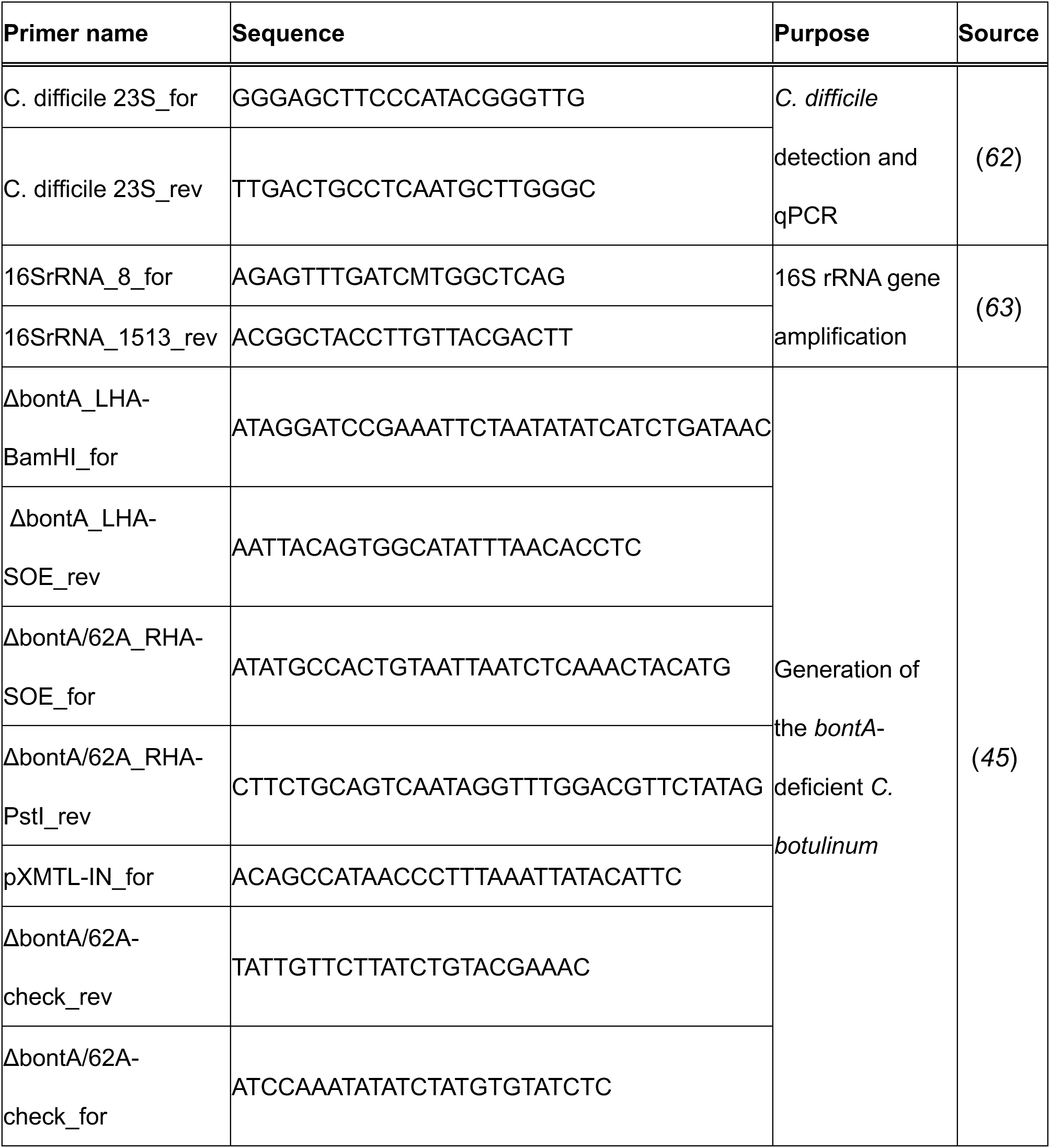
Primers used in this study.

**Table S7.**
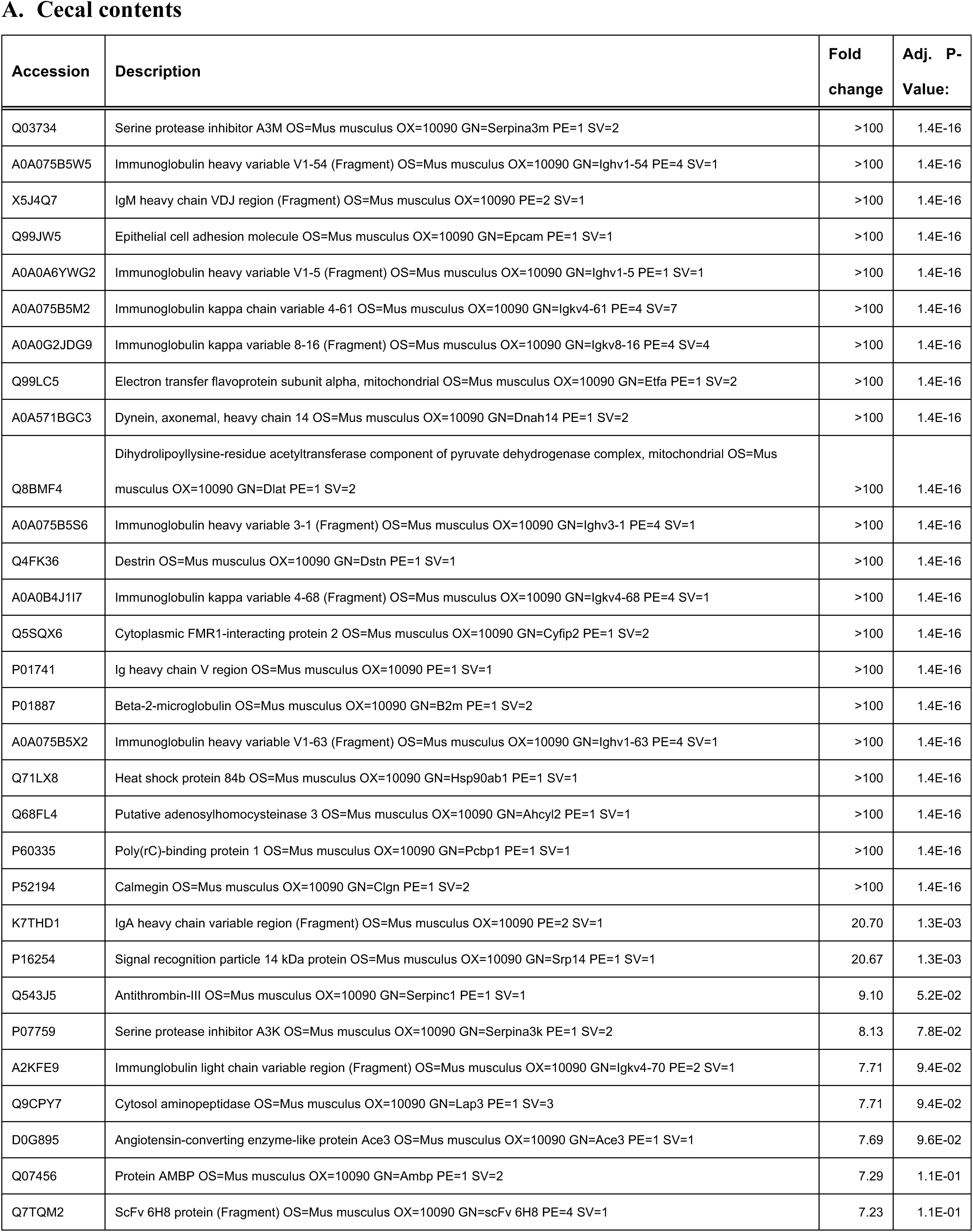
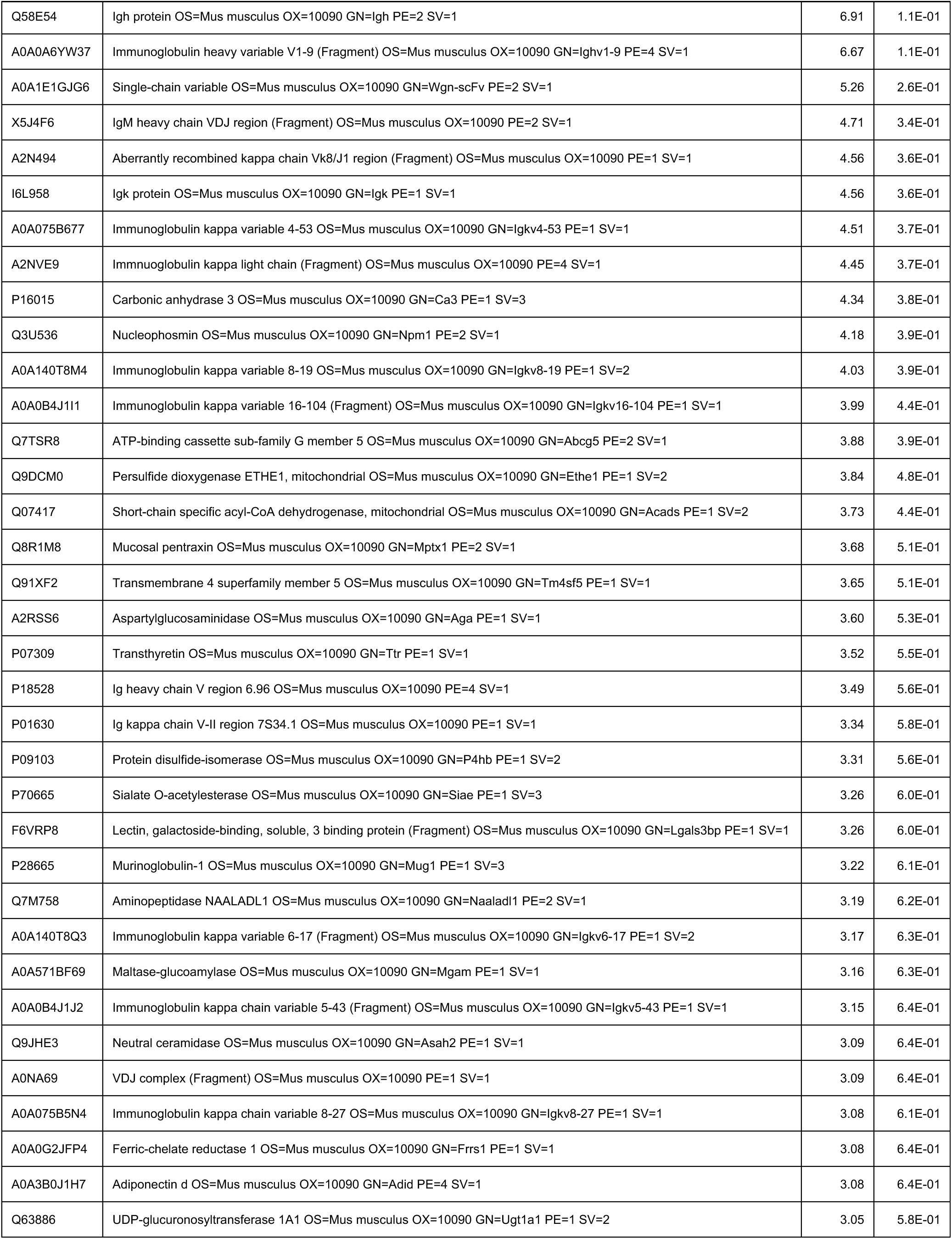
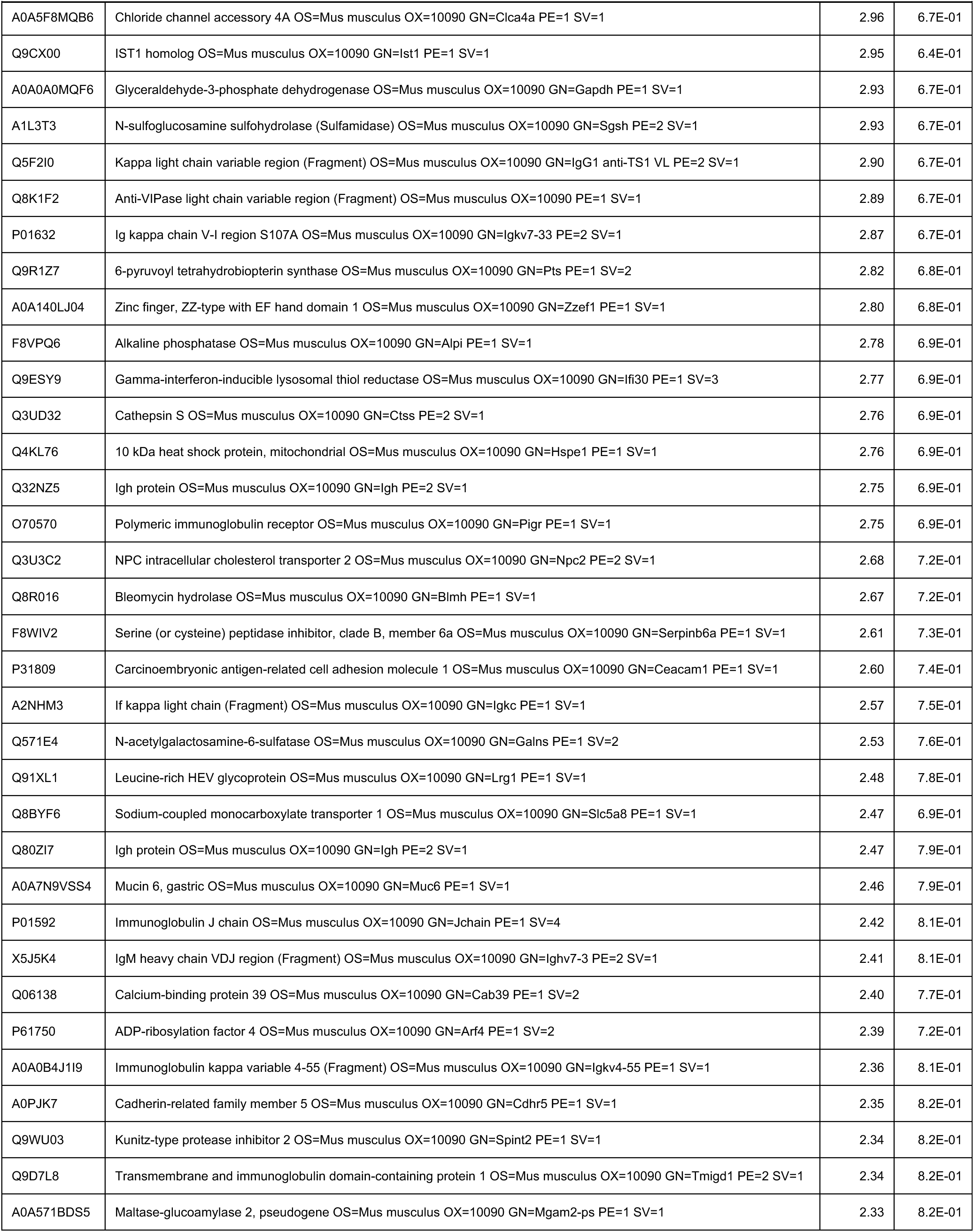
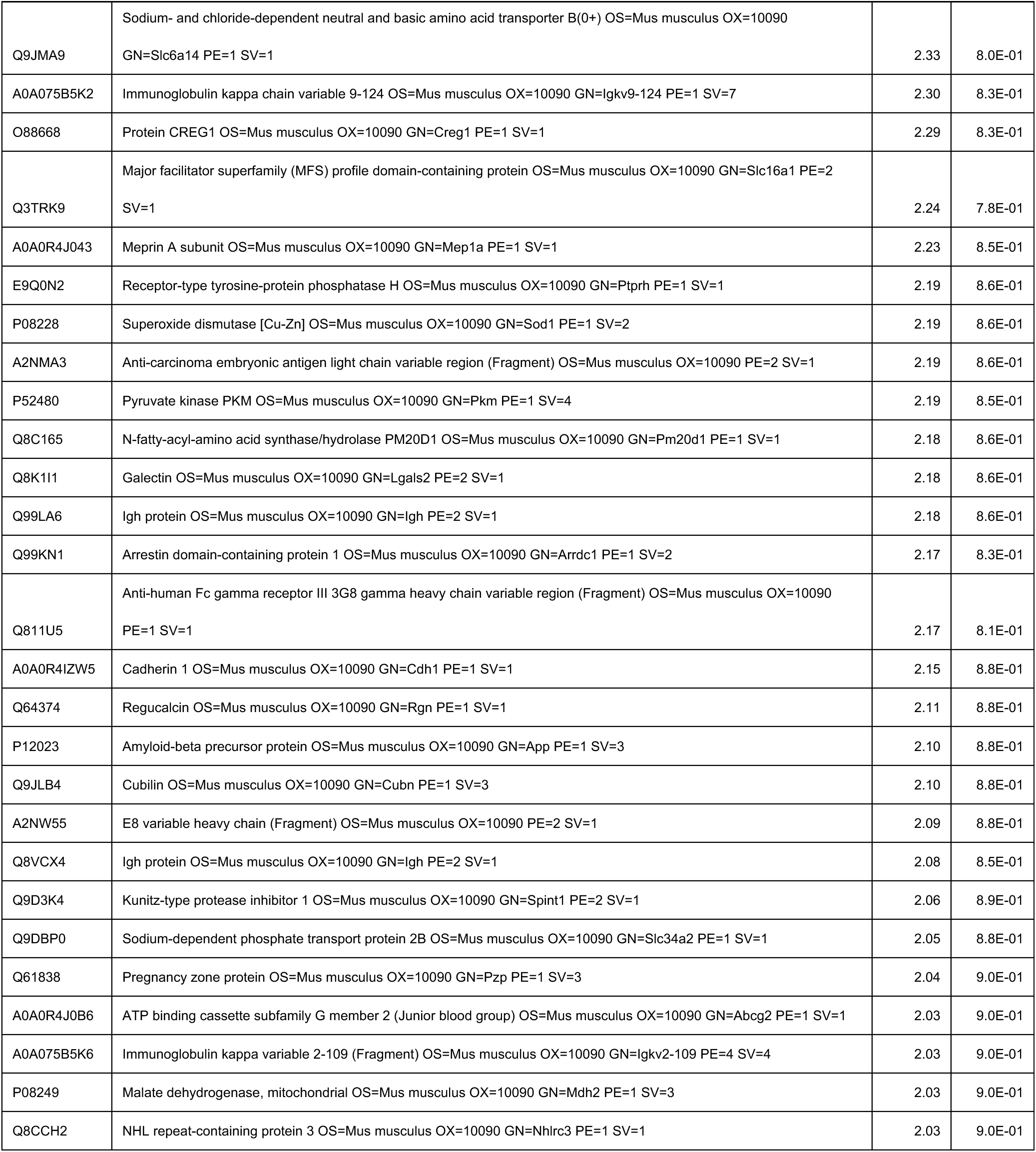
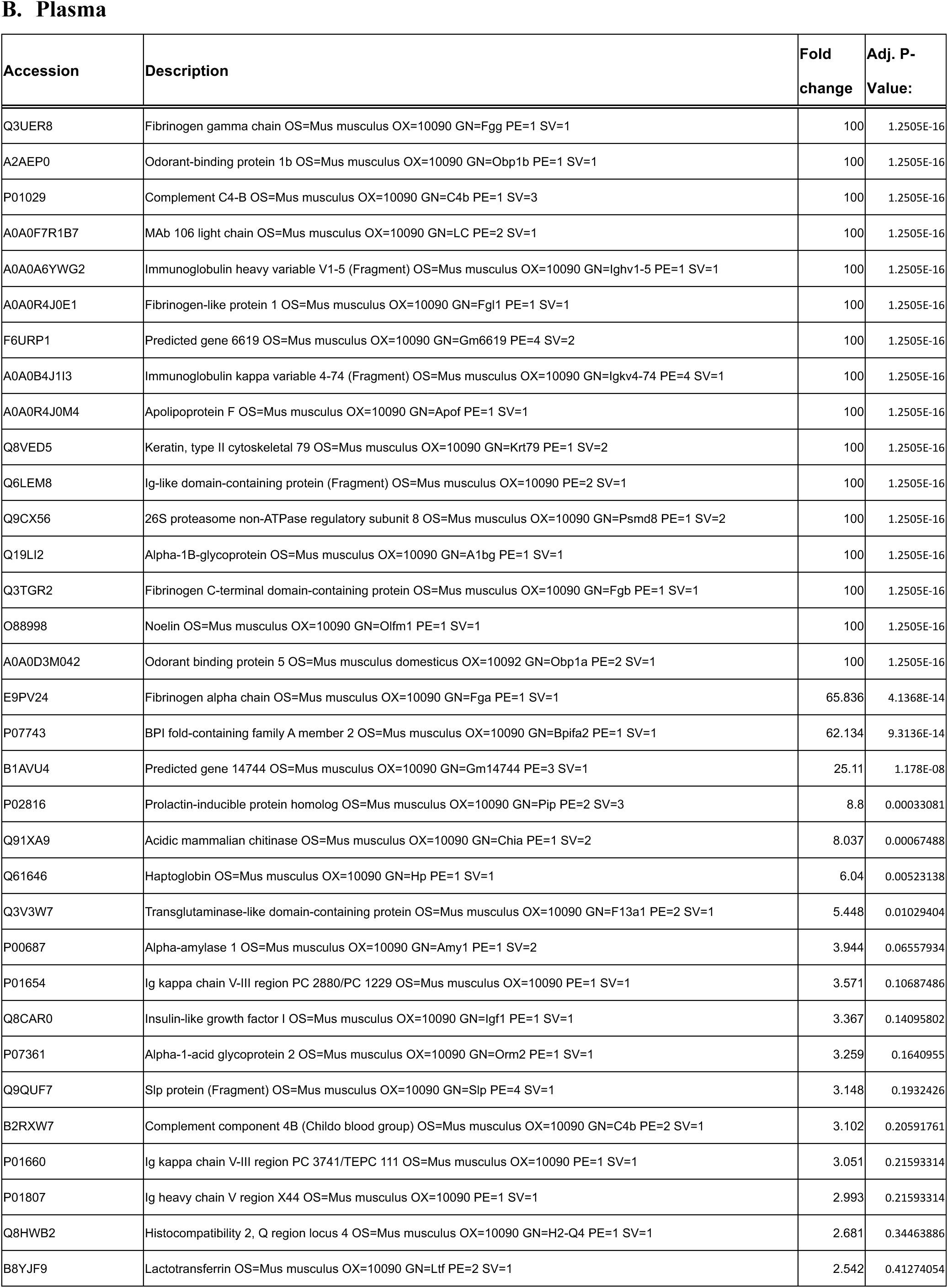
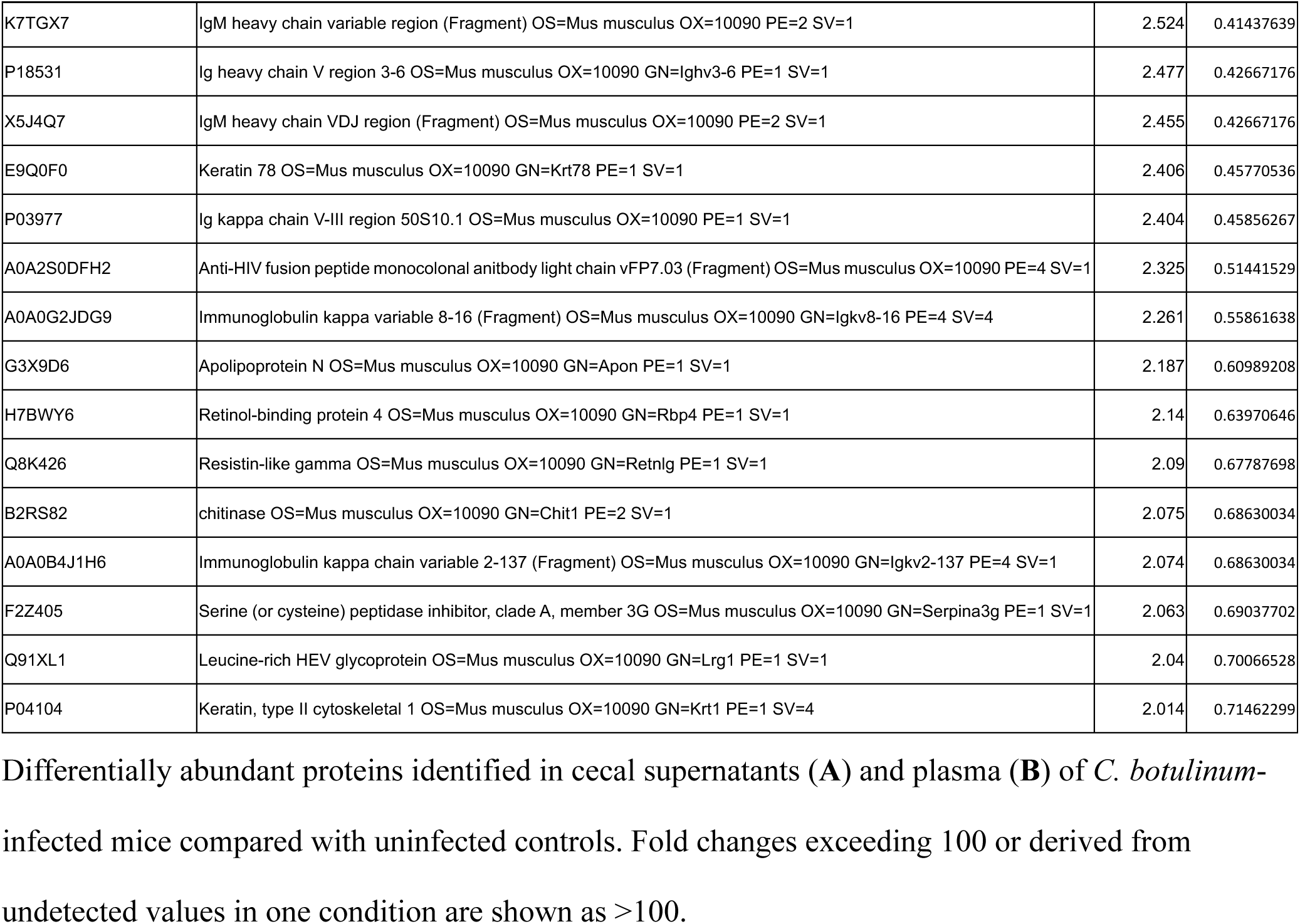
Proteins upregulated more than twofold *in C. botulinum*-infected mice identified by proteomic analysis.

## Notes

### Competing Interest Statement

Some authors are inventors on a pending Japanese patent application (No. 2026-112886) related to the findings reported in this manuscript. No payments or services from third parties that could be perceived to influence this work were received.

### Summary of Updates

Revised manuscript with improved organization, updated Discussion, and minor corrections to text, figures, and supplementary materials.

